# Main Manuscript for Microbiome single cell atlases generated with a benchtop instrument

**DOI:** 10.1101/2023.08.08.551713

**Authors:** Xiangpeng Li, Linfeng Xu, Benjamin Demaree, Cecilia Noecker, Jordan E. Bisanz, Daniel W. Weisgerber, Cyrus Modavi, Peter J. Turnbaugh, Adam R. Abate

## Abstract

Single cell sequencing is useful for resolving complex systems into their composite cell types and computationally mining them for unique features that are masked in pooled sequencing. However, while commercial instruments have made single cell analysis widespread for mammalian cells, analogous tools for microbes are limited. Here, we present EASi-seq (Easily Accessible Single microbe sequencing). By adapting the single cell workflow of the commercial Mission Bio Tapestri instrument, this method allows for efficient sequencing of individual microbes’ genomes. EASi-seq allows thousands of microbes to be sequenced per run and, as we show, can generate detailed atlases of human and environmental microbiomes. The ability to capture large shotgun genome datasets from thousands of single microbes provides new opportunities in discovering and analyzing species subpopulations. To facilitate this, we develop a companion bioinformatic pipeline that clusters genome by sequence similarity, improving whole genome assembly, strain identification, taxonomic classification, and gene annotation. In addition, we demonstrate integration of metagenomic contigs with the EASi-seq datasets to reduce capture bias and increase coverage. Overall, EASi-seq enables high quality single cell genomic data for microbiome samples using an accessible workflow that can be run on a commercially available platform.

## Introduction

A microbiome comprises the collection of distinct microorganisms and their genomic elements within a particular environment. These microecosystems play fundamental roles in the biosphere, have major impacts on human health, and are important resources for scientific and economic progress(O’Connor, 2021; Timmis et al., 2017). Thus, the study of microbiomes – in terms of the species present, the genes employed by different members to thrive, and the molecules consumed or produced – is of high scientific value. Historical methods of research rely on passive observation with microscopy(Blaser et al., 2016), which predominantly yield information about phenotypes and behavior. To reveal functional properties, assays can be conducted on microbes isolated from the environment(Brehm-Stecher and Johnson, 2004; Caumette et al., 2015); however, the need to culture microbes isolated from environment form many of these studies imposes a significant bias or precludes any detection(Kapinusova et al., 2023). Thus, culture-independent methods to profile microbes’ taxonomic identities and functions are immensely valuable. Amplicon sequencing of 16S ribosomal RNA (rRNA) genes or other diagnostic marker genes has been widely used for classifying microbial community composition. Despite its convenience, amplicon sequencing suffers from PCR bias and can have limited resolution in discriminating closely related species or strains of the same species(Jones et al., 2022). Metagenomic sequencing, which directly sequences all genomic DNA within an environment, enables both the profiling of phylogenetic diversity and the comprehensive accounting of all the genes present within a microbiome(Eloe-Fadrosh et al., 2016; Quince et al., 2017). However, because the data is acquired as a pool of mixed sequencing reads originating from all organisms, the bioinformatic reassembly requires sophisticated computational algorithms for assembly and sometimes yields disconnected genomic fragments and creates chimeric sequences of different strains or species (Meyer et al., 2022; Wooley and Ye, 2010). Although long read sequencing(Moss et al., 2020) and/or read cloud algorithms(Bishara et al., 2018) can generate relatively long genomic assemblies, associating separate chromosomes or chromosomes and extrachromosomal elements (e.g., plasmids) with a single cell type can still be challenging with current methods. Characterizing these associations can be critical to understanding the behavior of a microbiome and genetic flow. While approaches like genome-resolved metagenomics(Zhou et al., 2021) and chromosome conformation capture(Yaffe and Relman, 2020) can obtain them in some circumstances, these methods are biased towards the more abundant species/extrachromosomal elements and may generate spurious links between DNA fragment from different cells(Stalder et al., 2019)..

A microbiome consists of heterogeneous microbial populations(Berg et al., 2020). Thus, just as single cell sequencing has transformed mammalian cell biology by resolving heterogeneous systems and tissues into their composite cell types(Marioni and Arendt, 2017; Stuart and Satija, 2019), similar impacts are possible in microbiology. Of the possible methods, single cell genomics is perhaps most important for microbiology because of the significant genetic heterogeneity and frequent transfer of genetic material(Van Rossum et al., 2020). Genetic mobile elements can be a source of important phenotypes, including virulence factors or resistance genes that can transform a normally harmless commensal into a multidrug resistant pathogen(Frost et al., 2005; Soucy et al., 2015). Methods to analyze all genomic information of a cell, including DNA not physically connected to the chromosome, would allow characterization of these vital mobile elements. Towards this objective, there has been significant effort to develop single microbe sequencing. Previous approaches have been based on isolating microbes for single cell genome amplification and library preparation using FACS(Arikawa et al., 2021; Chijiiwa et al., 2020; Freiherr von Boeselager et al., 2018; Hosokawa et al., 2017; Imdahl et al., 2020; Lawrence et al., 2022; Nishikawa et al., 2022; Pachiadaki et al., 2019; Zhang et al., 2006), optical tweezers(Xu et al., 2020), hydrogel matrix embedding(Xu et al., 2016), and microfluidics(Marcy et al., 2007). These methods either have limited throughput, allowing just hundreds of genomes to be sequenced, or are highly labor intensive. More recently, barcoding reminiscent to scalable mammalian cell methods have been applied to microbes and achieved the sequencing of similar numbers of cells (thousands of cells per run)(Lan et al., 2017; Zheng et al., 2022). These multi-step droplet microfluidic approaches utilize robust molecular biology, yielding superb data for most cell types in the sample(Lan et al., 2017; Zheng et al., 2022); unfortunately, the number of steps and custom-built instrumentation poses a significant barrier to non-microfluidic engineers for its application. Another single step droplet microfluidic approach enables sequencing multiple gene loci of thousands of single bacteria cells were reported(Lan et al., 2024, 2023), however, its application is limited to only a few amplicons. Meanwhile, high-throughput single bacteria RNA sequencing has been demonstrated using combinatorial indexing(Blattman et al., n.d.; Kuchina et al., 2021) and commercially available single cell platforms(Ma et al., 2023). However, these methods have only been used for model organisms and have never been applied to a complex microbiome, in which the diverse physical properties of microbes make optimization of the requisite fixation, permeabilization, and in situ ligation difficult. Thus, currently, there is no high-throughput method available to the microbiological community for efficient single cell genome sequencing of microbiomes. If such a method could be developed, it would be superior to metagenomic sequencing in most instances and provide access to capabilities currently missed, including generation of complete single-microbe resolution cell atlases and gene annotation at the strain or single cell level.

In this paper, we describe EASi-seq (Easily Accessible Single microbe sequencing), a method to efficiently sequence tens of thousands of microbes. Rather than relying on custom microfluidic instrumentation as in previous methods(Lan et al., 2017; Zheng et al., 2022), we start from a commercially available workflow with the inherent capabilities for single cell sequencing: Mission Bio’s Tapestri(Xu et al., 2019). This instrument is widespread in clinical and academic centers, easy to use, and reliable. A major impediment is that the instrument is designed for targeted DNA sequencing of mammalian cells and is not directly applicable to microbial cell whole genome sequencing, which requires different lysis strategies and nucleic acid adaptors. To address this, we introduce two key modifications into the commercial workflow: a bulk single cell nucleic acid purification step that addresses cell lysis and adapter tagmentation(Picelli et al., 2014) protocol that enables whole genome sequencing. With these modifications, the Tapestri generates barcoded sequence data for several thousand microbial cells which, as we show, can comprise bacteria, archaea, and fungal linages. The data generated by EASi-seq are untargeted single cell shotgun reads across each cell’s genome and any other DNA present in the cell, such as mobile elements and plasmids. To facilitate analysis of the single cell sequencing data, we develop a companion bioinformatic pipeline that clusters cells into similarity groups, annotates their genes and species, and pools sequences within a cluster to increase improve genome assembly and coverage. Using EASi-seq, we generate detailed atlases of a control synthetic community and human gut microbiome and costal water microbiome. We show that EASi-seq’s single cell resolution allows differentiation of microbial strains with 99% genomic similarity. EASi-seq provides a universal approach for deconvoluting microbiomes into the cells of which they are composed and to characterize their gene and pathway functions.

## Results

### EASi-seq workflow for whole genome microbial sequencing

A platform to reliably sequence large numbers of environmental microbes must overcome technical and practical challenges. Different microbes have different cell wall and membrane properties and, thus, can require a mixture of different lysis reagents or procedures(Lan et al., 2017; Maghini et al., 2021; Yuan et al., 2012). Additionally, genomic and plasmid DNA must be fragmented and have the correct adaptors added prior to being sequenced. Lastly, some microbiomes are highly heterogeneous, having hundreds to thousands of distinct species that potentially include many strains. Generating a complete single cell atlas in this scenario requires sequencing significant numbers of single cells. Prior methods to overcome these challenges used custom workflows with 3 to 5 microfluidic processing steps(Lan et al., 2017; Zheng et al., 2022). Each device had to be custom fabricated and operated by microfluidic experts. While the works demonstrate the power of high throughput single microbe sequencing, the inaccessibility of these workflows precludes their use by microbiologists lacking microfluidic expertise. Recently, several commercial single cell instruments have become available that support processes like the ones required for microbial whole genome sequencing (**Table S1**). Of these, Mission Bio’s Tapestri is unique in the ability to conduct two subsequent droplet steps as a result of being designed for targeted DNA sequencing of mammalian cells(Xu et al., 2019). Nevertheless, even with automation of two common microfluidic steps, directly replicating prior microbe sequencing workflows on Tapestri is not possible. Thus, a major innovation of this work is to develop a microbe sequencing workflow that maps onto Tapestri’s two-step process.

To enable single microbe sequencing, the cell must be lysed, the DNA fragmented into readable lengths, and the fragments labeled with single cell barcodes. With Tapestri’s two step workflow, we can use the first droplet manipulation stage to perform DNA tagmentation, and the second for barcoding. The challenge is lysing the cells to prepare the genomes for tagmentation, while keeping the genomes and extrachromosomal DNA together. A proven approach to accomplish this is to use a microfluidic device to encapsulate single cells in hydrogel spheres before lysing the cells with bulk washes. Because genomic DNA is a high molecular weight polymer, it remains ensnared within the hydrogel matrix and is protected from the shear forces generated by washing, allowing intact genome purification(Arikawa et al., 2021; Chijiiwa et al., 2020; Hosokawa et al., 2017; Lan et al., 2017; Nishikawa et al., 2022; Spencer et al., 2016). The requirement of microfluidics for cell encapsulation, however, would negate the primary advantage of EASi-seq’s accessibility. Thus, another core element of this work has been to develop a microfluidic-free process for genome purification in hydrogels. In the approach, we encapsulate the cells in hydrogel droplets by emulsification through vigorously shaking or shearing through a syringe needle (**Fig. 1a**). In either case, the process generates an emulsion in which the cells are randomly loaded. The droplets of polyacrylamide monomer are gelled by radical polymerization to ensnare the cells. The resulting hydrogel beads are then transferred into an aqueous carrier for lysis and washing. This process takes 2 hours and uses no microfluidics. The approach uses radical polymerization reactions that have been shown to not damage DNA damage and be compatible with PCR and genomic sequencing(Spencer et al., 2016). The resultant suspension is polydisperse, containing many hydrogel beads too large for the Tapestri (which only accepts cells or beads having a diameter less than 30 µm) or too small to trap a cell. Thus, we use differential centrifugation (**Fig. 1b** and **Fig. S1**) to select hydrogel beads sizes within the optimal 5-30 µm diameter range. To ensure a high probability of single cell genomes, the initial cell concentration is set such that hydrogels of this size are loaded at a rate of 2%. To purify the genomes, we perfuse the hydrogel beads with cocktails comprising polysaccharide digesting hydrolases and proteolytic enzymes(Lan et al., 2017). The result is a suspension of hydrogel beads with intact single cell genomes that have similar physical and hydrodynamic properties to mammalian cells. These beads can be readily processed with the Tapestri (**Figs. 1c-d**). To ensure single cell sequence data, most of the hydrogels are left empty, such that about 10% contain single cells, in accordance with Poisson statistics(Collins et al., 2015). Thus, when loading the gels into Tapestri, we set the concentration to about 5 gels per droplet, which yields 10% containing one genome and 90% containing no cells, thereby yielding single cell data, and making efficient usage of the barcoding droplets.

**Figure 1.**
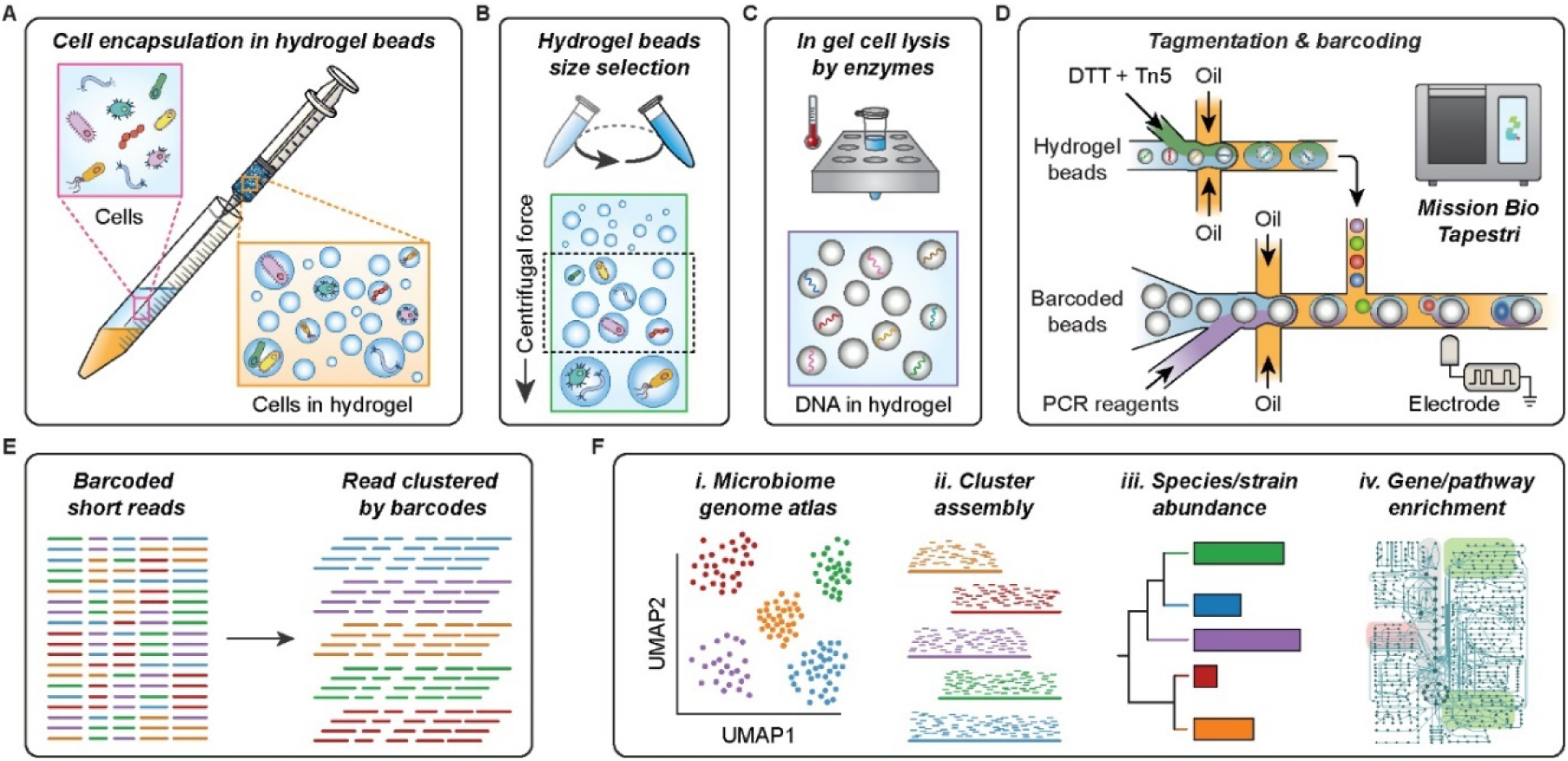
EASi-seq genome purification, microfluidic, and bioinformatic workflow. (**A**) Microbial cells suspended in hydrogel precursor (acrylamide monomer and BAC crosslinker) are emulsified with a fluorinated oil by passing the mixture through a syringe needle. After gelation, the cells are individually embedded in hydrogel beads. (**B**) The hydrogel beads are size selected using differential centrifugation. The hydrogel beads are suspended in a density matching buffer (40% sucrose in PBS with 0.1% Tween 20) and centrifuged. Particles of different size sediment at different rates, with the larger particles sedimenting faster. After centrifuge at 1000 x g for 10 min, the oversized hydrogel beads are pelted, and the supernatants are subject to a centrifuge at a higher speed (3000xg for 10 min). The pellets are then collected as the size-selected hydrogel beads. (**C**) Cells are lysed within the hydrogel beads by a two-step enzyme digestion. The beads are first subjected to a cocktail of 4 different enzymes that digest cell walls before being treated with protease K to digest proteins. The small pore-size of the hydrogel allow proteins and other molecules to freely diffuse, while immobilizing long DNA molecules. After the treatments and washing, only genomic DNA remains in the hydrogel beads. (**D**) The microbial genomic DNA in each hydrogel bead is tagmented in a droplet (first step, *bottom*) before being subsequently paired with barcode beads for barcoding PCR (second step, *top*) on using the Tapestri instrument’s microfluidic modules. (**E**) Sequencing of amplicons from the barcoding PCR generates single-cell shotgun reads for thousands of cells. (**F**) EASi-seq allows high-throughput microbiome genome atlas analysis, as well as cluster-based genome assembly, strain identification, and pathway analysis.

Normally, the Tapestri’s first step is used to encapsulate and lyse the cells. Since our cells are already lysed in the gels, we can use this module for tagmentation instead. To maximize tagmentation efficiency, the genomes must be released from the hydrogel beads. Controlled release is accomplish by utilizing N,N’-bis(acryloyl)cystamine (BAC) as the hydrogel crosslinker, which can be reversed on-demand with dithiothreitol (DTT) addition(Wang et al., 2020). The Tapestri’s dual-inlet design for the first step allows DTT addition with Tn5 transposase, such that the hydrogels liquify upon droplet encapsulation (**Fig. 1d, *top module*** and **Fig. S2**). The Tn5 transposase used for tagmentation is loaded with forward adaptors matching Tapestri’s V2 barcoding primer 3’ constant region (**Table S2**), allowing the tagmented fragments to be barcoded in the subsequent droplet PCR (**Fig. S3**). At this point, genomic DNA is released and tagmented in each newly formed droplet. Barcoding is accomplished by droplet reinjection and merging with the needed barcoding PCR reagents in the Tapestri’s second step (**Fig. 1d, *bottom module***). After the barcoding PCR, the final sequencing adaptors are added by pooling the amplicons of all droplets and using a bulk PCR (**Fig. S3**). The resultant material is sequenced and computationally deconvoluted into single cells by barcode (**Fig. 1e**). The datasets contain tens of thousands of single cell genomes with coverages ranging 0.01-10% depending on genome size and sequencing depth. The genomes can be clustered into a single cell atlas (**Fig. 1f, *i**.-ii.***). The data for all single cells in each cluster can be pooled to create a consensus genome. Metagenomic sequencing data can be integrated to increase coverage. The final genus clusters can be annotated and evaluated for features of interest, including species or strain abundance and gene or pathway distributions (**Fig. 1f, *iii.-iv.***).

### Validation of single cell resolution

For EASi-seq to be useful, it must generate barcoded single cell sequence reads. To validate this capability, we used EASi-seq to analyze the synthetic ZymoBiomics microbial community, consisting of eight bacteria and two yeasts (**Fig. 2a**, **Table S3**). We process the community using EASi-seq, generating 205,730,606 paired-end reads in 14,175 barcode groups after filtering by barcode read counts (**Fig. 2b)**. The barcode groups showed two distinct populations, one (with 10,543 barcode groups) had low alignment rate (2.15%), while the other had high alignment rate (91.61%) (**Fig. 2c**). We reason that the low alignment barcode groups are the non-specific PCR products in the droplets without any cells. In absence of cells, the oligonucleotides assembled on the Tn5 enzyme might interact with the barcode primers and generate amplicons. However, when cells are present, the tagmentation products are dominantly amplified. Thus, we remove the barcode groups with low alignment rates (**Fig. 2c**) and recover 3632 barcode groups with an average of 38, 576 reads (ranging from 2000 to 1,931,407 reads). To assess single cell resolution, we map the reads in each group to the ten reference genomes and plot the fraction mapping to the dominant species. We find that 86.16% of barcodes have a purity of >90% (**Fig. 2d**) and dominantly align to one species (**Fig. S4, Table S4**). These results demonstrates that EASi-seq achieves single cell resolution. For the shallow sequencing applied, the average coverage is 0.44% for bacterial genomes and 0.031% for the larger yeast genomes (**Fig. 2e**). The coverage of most single cell barcode groups remains unsaturated at 10,000 reads (**Fig. S5-6**). The comparison of EASi-seq with metagenomic sequencing indicates gram-negative bacteria are poorly represented within EASi-seq barcode groups (**Fig. 2f**). For example, we identify only four *P. aeruginosa* cells with a total read count of 34,021. This result is consistent with a previous report (Lan et al., 2022) and is caused by the ZymoBIOMICS synthetic community inactivating buffer (DNA/RNA Shield^TM^) pre-lysing gram-negative bacteria.

**Figure 2.**
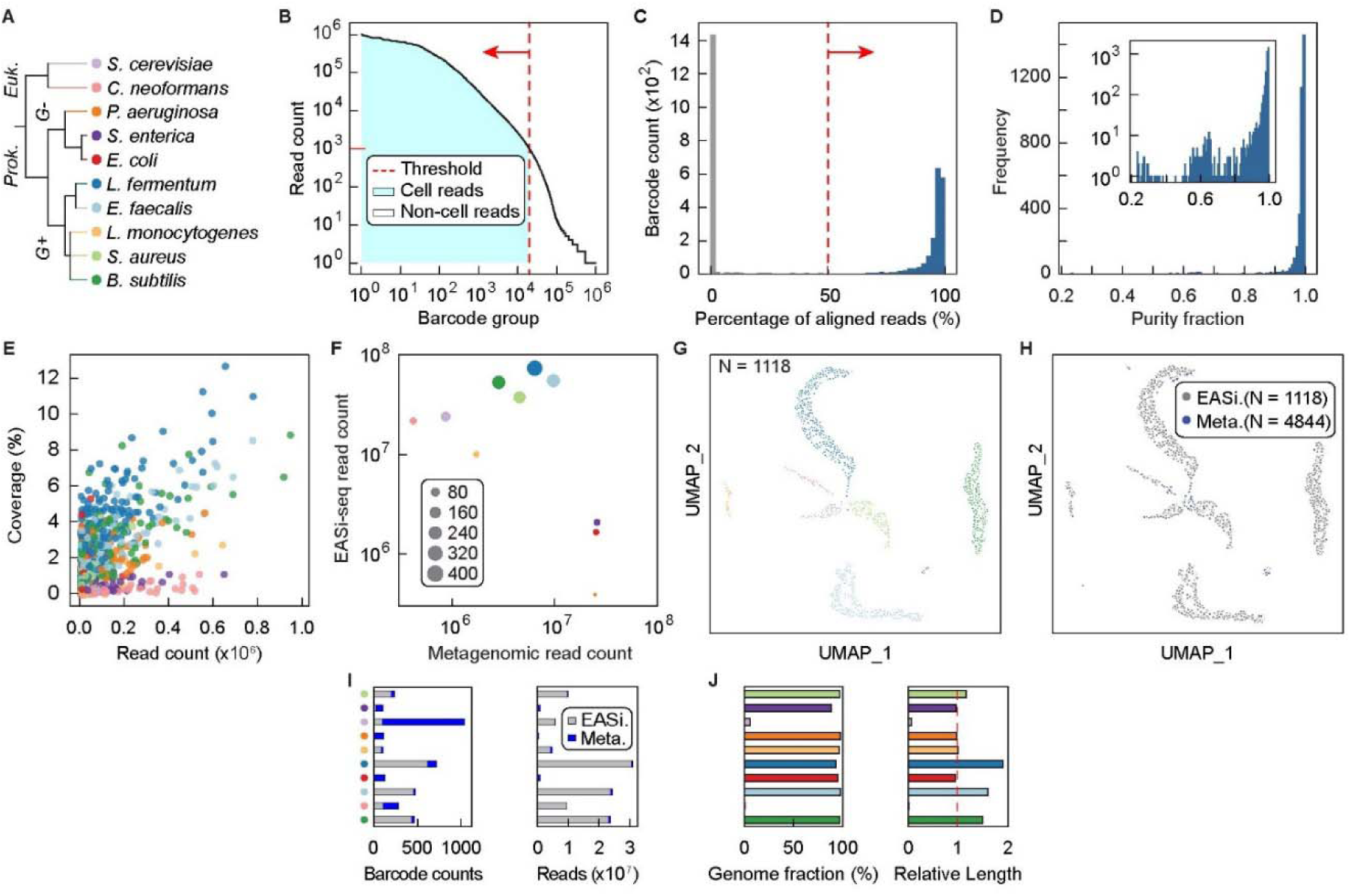
EASi-seq identifies single cells and has strain-level resolution. (**A**) The ZymoBIOMICS microbial synthetic community consisting of 10 species was analyzed by EASi-seq. Classification of each species is provided, with assigned colors used in the following panels. (**B**) Barcode rank plot of obtained data. Barcode groups were filtered by read counts, with less than 1000 reads used as the cutoff. (**C**) The barcode groups were further filtered by alignment rates to reference genomes. Barcode groups are mapped to the combined reference genomes of the 10 species, and barcode groups with an alignment rate of less than 50% were removed. (**D**) Purity distribution of barcode groups after data filtering, defined as the percentage of the reads mapping to the species most represented among read alignments. Inset shows purity distribution as a log-scale. (**E**) Coverages of each barcode group, color coded by species. (**F**) The comparison between metagenomic and single cell sequencing. Scatter plot shows the read counts of metagenomics sequencing data and combined EASi-seq barcode groups. Data points are color coded by species and their sizes are proportional to barcode counts after filtering. (**G**) UMAP clustering by Taxonomic discovery algorithm, color coded by species. Each barcode group is classified using a k-mer based taxonomy classifier (Kraken2). The output files were combined at the genus level. The barcodes were filtered by the percentage of mapped reads and taxonomical purity, which is the percentage of the dominant taxa. The vector of the genus abundance in each barcode was used to generate the UMAP and each barcode is annotated by the most abundant genus. (**H**) UMAP clustering shows the integration of the EASi-seq data (gray) and metagenomic data (blue). Each contig associated short read group in the assembled metagenome of the same sample was treated as a barcode and processed by the Taxonomic discovery algorithm. (**I**) Barcode counts and read counts in each UMAP cluster, grouped by batch (EASi-Seq or Metagenome assembly). (**J**) Evaluation of contigs assembled by grouping reads from all barcodes in each cluster. All the reads within a cluster were assembled into contigs using Spades and evaluated by Quast using the reference genome. *Left*, Genome coverage. *Right*, relative contig length normalized to reference genome.

### Reference-independent clustering of unknown cell types

When applying EASi-seq to a novel microbiome, reference genomes are usually not available for mapping and species assignment of the single cell datasets. Thus, to build a genome atlas that displays all cells in a sample, we require a clustering algorithm not reliant on prior knowledge of the species present. In addition, many single cell genomes are covered below 1% (**Fig. 2e**) and comprise short reads that do not overlap with other single cells of the same type in different barcode groups. To enable clustering from such data, we propose the Taxonomic Discovery Algorithm (TDA). In TDA, each barcode group is treated as a metagenomic sample, and its taxonomic abundance is estimated with available taxonomic classifiers. The taxonomic estimations of all barcode groups are then combined into a vector suitable for similarity clustering. We hypothesize that the different barcode groups that belong to the same cell should be classified to the same taxa by taxonomic classifiers even if they possess completely different sets of reads. In this approach, reads of each barcode groups are first classified based on a taxonomic database to estimate the barcode group’s associated taxonomy abundance. The taxonomic abundances of all barcode groups are binned into a vector consisting of all genera, wherein the bin value is proportional to the number of reads in the barcode group mapping to it based on the available taxonomic database. For taxa accurately represented in the database, most reads will be assigned to one genus bin, while cells from poorly represented taxa may be assigned to several. The core concept is that related cells generate similar genus vectors, even if their reads cover different portions of the genome, and even if the genus to which individual reads are assigned do not perfectly match the true genus (**Methods**). In addition, barcode groups with many nonspecific amplification products do not match any taxonomic groups and are thus removed. With the genus vectors in hand, related cells can be clustered using Uniform Manifold Approximation and Projection (UMAP)(Becht et al., 2019) for visualization. After clustering, reads from all cells in a cluster are pooled to generate a consensus genome.

The efficacy of the TDA for clustering cells depends on the classification method and database used to map reads into genus bins. If a species is totally novel, such that few of its reads can be annotated to any genera or other taxa level, the vector will contain minimal useful information for clustering. To identify the best database for the TDA, we therefore evaluated the most popular software for taxonomic classification and quantification. These included tools based on K-mer (Kraken2/Bracken(Lu et al., 2017; Wood et al., 2019)), marker gene (MetaPhlAn3(Beghini et al., 2021)), and protein similarity (Kaiju(Menzel et al., 2016)). For this evaluation, we simulated a microbiome using downloaded available genomes, processed into single cell barcode groups with read structures resembling the output of EASi-seq (**Methods, Fig. S7a-b**). To assess the efficacy of a classification method, we calculated the accuracy of barcode purity prediction and taxonomic annotation, and barcode recovery rate with filtering. The k-mer based Kraken2/Bracken with PlusPF database (v.2021/01/27)(“Kraken 2, KrakenUniq and Bracken indexes,” n.d.; Lu et al., 2022) showed the best performance in genus identification accuracy, purity prediction accuracy, and accurate barcode retention rate (**Discussion S1, Fig S7c-h**) and was our choice going forward.

With the TDA validated on a simulated dataset, we next experimentally verified it using the known composition of the ZymoBIOMICS synthetic community. After filtering based on the percentage of mapped reads to remove the non-specific amplicons and based on genus level purity to select the pure barcode groups (**Fig. S8a-b**), the TDA correctly clustered and identified all ten populations in this synthetic community (**Fig. 2g**, **Table S5**). In addition, 97.34% of barcode groups were correctly annotated, reflecting the good representation of these community members in that database (**Fig S8c**).

### Integrating metagenomic contigs to reduce cell-type bias and increase overall coverage

A unique and powerful feature of EASi-seq when combined with unbiased clustering is the ability to pool single cell data to increase genome coverage. Compared to EASi-seq, metagenomic sequencing does not rely on intact cells, and uses all extracted nucleic acid that may better capture all microbial taxa. Thus, to enhance the coverage of EASi-seq, we developed an approach to integrate metagenomic data using a similar strategy to the TDA, in which we calculate a genera abundance vector for each contig assembled from metagenomics data, then co-cluster the metagenomic contigs with the single cell barcode groups (**Fig. S9**, **Table S6, Methods**). These vectors are filtered by purity (**Fig. S10**) before clustering. From a metagenomic assembly of the ZymoBiomics community, we identified 1427 of 4844 contigs that had >90% association with one genus. Most contigs clustered in a fashion that overlapped with the single cell data points (**Fig. 2h, Fig. S11**). With reads added by metagenomic contig integration, we achieve an average cluster coverage of 94.31±4.92% for bacteria and 2.74±3.24% for fungi, and the relative contig lengths approach 100% of the genome (**Fig. 2i-j**). Additionally, the assembled contigs have a GC content consistent with the reference genomes (**Fig. S12a**) and an average N50 of 49 Kbp (**Fig. S12b**). These results demonstrate that integration of metagenomic contigs with the EASi-seq atlas enhances capture of diverse microbial taxa and increases genome coverage.

### Strain-resolved differentiation

Differentiating between strains within a species is important for analysis of natural and engineered microbiomes(Van Rossum et al., 2020). Because EASi-seq can obtain thousands of reads on each cell, it affords novel opportunities for strain differentiation. To evaluate the ability of EASi-seq to accomplish this, we used it to analyze a synthetic community consisting of twenty-two equally mixed strains of *Eggerthella lenta*(Bisanz et al., 2020) (**Fig. 3a, Table S7**). We sequenced the library at 105,896,184 paired-end reads after quality filtering. We grouped the reads by barcode and aligned them against the reference genomes. To ensure read quality, we filtered barcode groups based on read counts and alignment rate (**Fig. S13**), recovering 5345 barcodes containing 101,760,151 reads. Because the strains have highly overlapping genomes, most reads align to multiple strains; thus, only reads specific to a single genome are useful for strain identification. Based on this, we developed a strain resolution approach reminiscent of transcript isoform expression estimation (BitSeq)(Glaus et al., 2012). We treat each genome as an isoform of one gene and estimate their “expression” level in each barcode group using BitSeq (parseAlignment and estimateVBExpression functions). All reads in a barcode group are mapped to the isoforms/strains and the probabilities of reads originating from a given isoform/strain are calculated for each alignment using a sequence-specific bias correction method (parseAlignment). Alignment probabilities are then used to calculate the posterior distributions of each isoform/strain via variational Bayes inference (estimateVBExpression), which is used to determine which strain a given cell most closely resembles (**Methods**). We aligned the reads in each barcode group to the reference genomes and recorded the overlap, using a Log-Normal read distribution to calculate the probability of originating from each reference genome, accounting for quality scores and mismatches. The barcode group is then assigned to a strain with more than 15% abundance and the highest abundance. (**Fig. S14**, **Table S8**). To visualize the resultant annotations, we plot the data as a UMAP and pair plot of the abundance estimation (**Fig. 3b and Fig. S15**). The separation between clusters on the UMAP plot confirms EASi-seq’s strain-level resolution.

**Figure 3.**
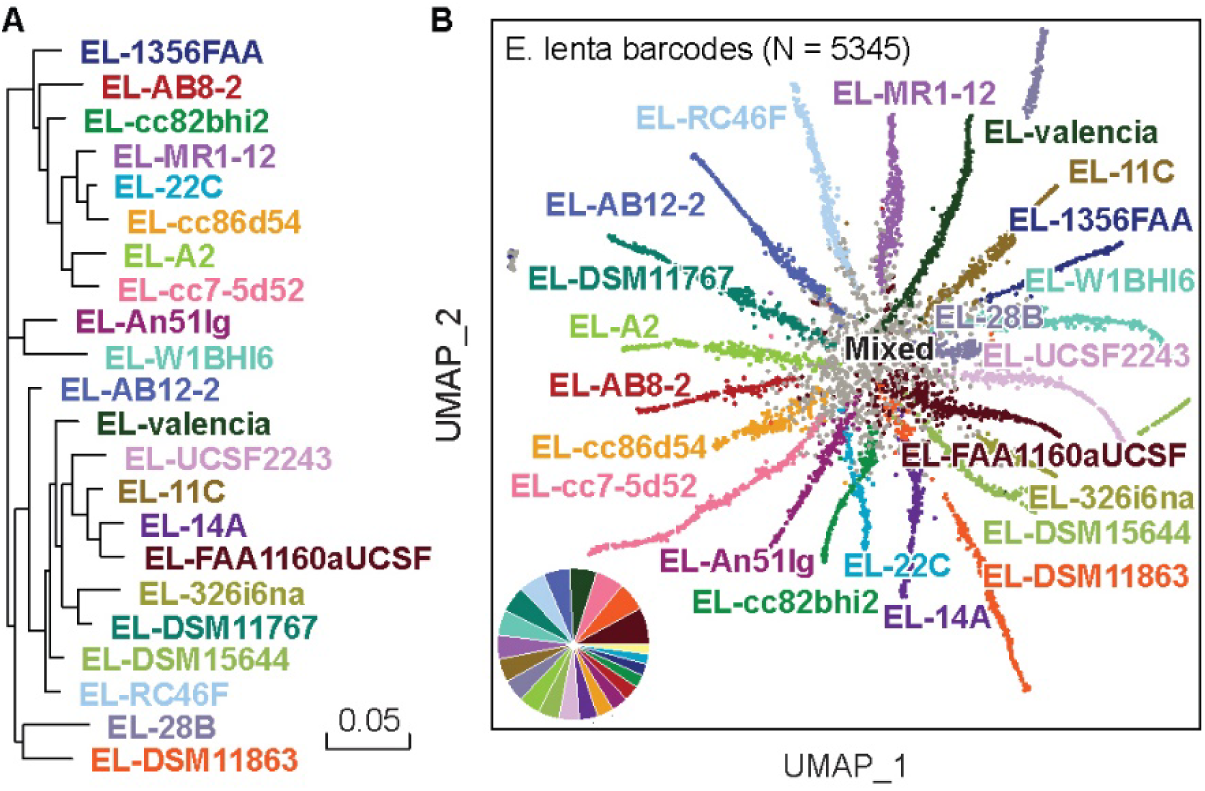
Strain resolution discrimination of microbes is achieved by EASi-seq. (**A**) Phylogenetic tree of the twenty-two *E. lenta* strains that make up the synthetic community. (**B**) UMAP clustering based on Bayesian abundance estimation of strains in each barcode group. Colors are the same as in (**A**), with any mixed/unresolved barcodes colored grey. The inset pie graph quantifies the fraction of barcode counts corresponding to each strain (excluding the mixed or unresolved barcodes), showing agreement with the expected equal distribution of strains.

### Single cell atlas of a human gut microbiome

The human gut microbiome comprises vast numbers of microbes from hundreds to thousands of species(Leviatan et al., 2022). Additionally, it can vary between individuals as a result of time, diet, geographical location, and health(Gilbert et al., 2018). Thus, characterizing microbiomes, the microbial taxa present, and their genetic functions is critical to understanding the dynamics and complexity of this ecosystem. Most approaches use amplicon sequencing of the 16S rRNA gene or bulk metagenomic sequencing(Wensel et al., 2022; Ye et al., 2019). EASi-seq would provide unique information missed by these methods, including single cell-level heterogeneity and cell-cell interactions. To explore this possibility, cells isolated (Hevia et al., 2015) from the human gut microbiome of a healthy donor were profiled by EASi-seq (**Fig. S16a**). After quality filtering, we recovered 232,705,096 paired end reads. We grouped reads by barcode and filtered by read count (>1000 reads) and genus purity estimated by Kraken2 (>80%) to remove multiplets and cell aggregates (**Fig. S17a-b, Discussion S2, Table S9**). The recovered 1118 barcode groups contained ∼150,000 reads on average. To increase cell capture efficiency and genome coverage, we also performed metagenomic sequencing of the sample (**Table S10**) and integrated it into the single cell data as described previously (**Fig. S9**, **Fig. S18a-c**). We filtered contigs based on read percentage classified by Kraken2 and genus level purity before integration with the EASi-seq data (**Figs. S18d-f**). We generated a cell atlas, identifying 95 clusters or microbial populations (**Fig. 4a**) with varied cell numbers and read counts (**Fig. S19)**. The metagenomic data increased the number of unique reads and clustered well with the single cell data (**Fig. S21, Table S11**). Nevertheless, several genera remain underrepresented in the atlas, including *Bacteroides*, *Phocaeicola*, *Parabacteroides*, *Akkermansia*, *and Alistipes* which may be a result of the cell isolation(Hevia et al., 2015) or sample storage artifacts(Watson et al., 2019), as has been described previously (**Fig. S20**, **Discussion S3**).

**Figure 4.**
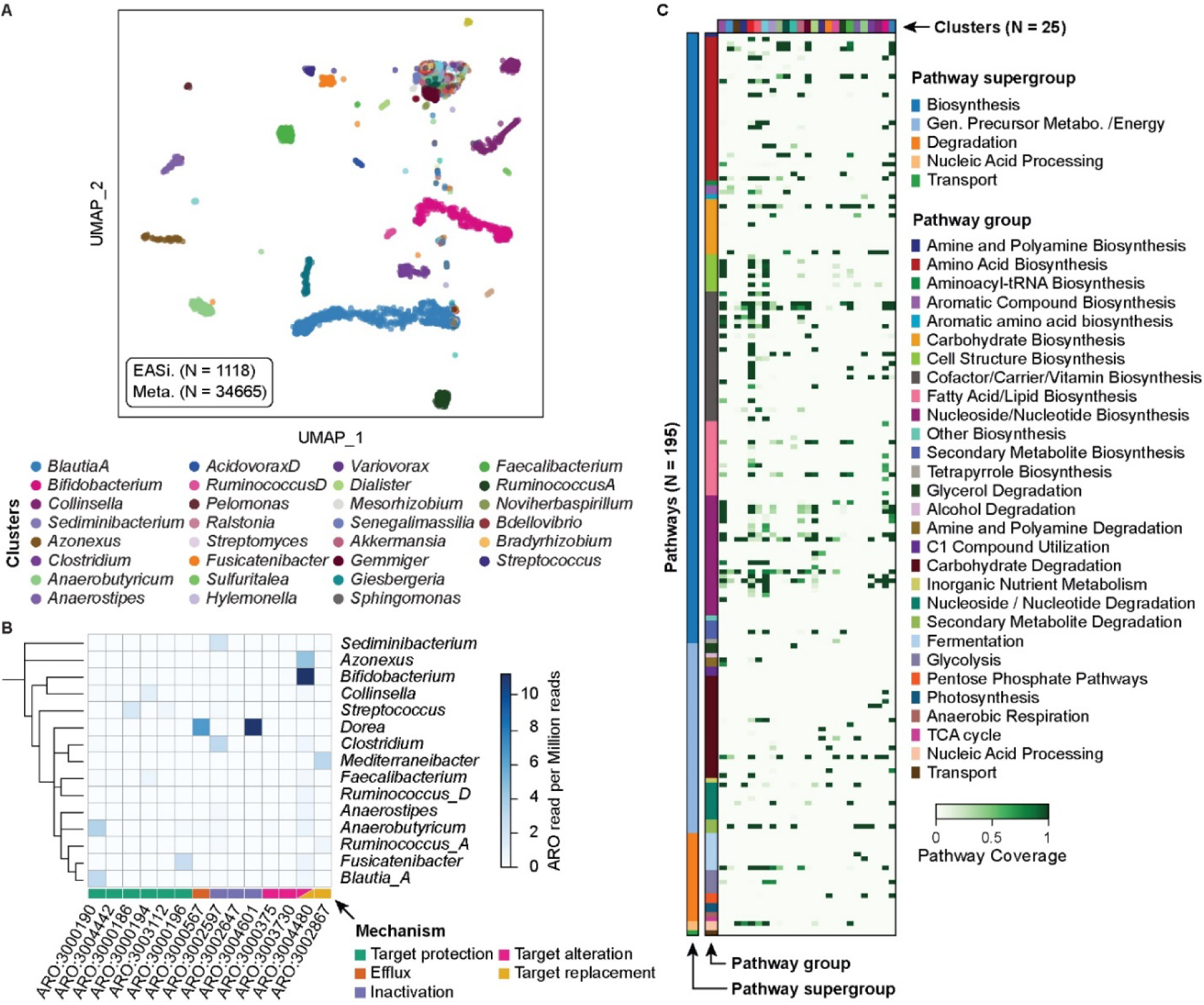
Human gut microbiome genome atlas. (**A**) Integrated UMAP clustering of the single cell barcodes and metagenomic assembled contigs of a human microbiome sample. Each barcode/contig was annotated based on the most abundant genus. Only the top 30 clusters are labeled in the legend. (**B**) Antibiotic resistance gene distribution in the clusters identified by TDA. Rows represented annotated genus and columns represent resistance gene access numbers from the Comprehensive Antibiotic Resistance Database. The value represents the read counts of the corresponding antibiotic resistance gene per million combined reads of each cluster. (**C**) Relative pathway abundance in the identified clusters. All reads in EASi-seq barcode groups associated to each cluster were combined and analyzed using MetaPhlAn and the MetaCyc database. The relative abundances of each pathway (copy per million, CPMs) were normalized to the barcode counts in each cluster. Clusters are color coded to the genera listed in (**A**).

The taxonomic level of the clustering depends on the taxonomic level used for the mapping in the TDA. Since we used genus for the analysis so far, clusters in the UMAP most closely represent this level. Thus, some clusters may group cells from multiple species, which may be resolvable by isolating these groups and re-clustering with a TDA analysis that uses species-level Kraken2 estimation (**Fig. S22**). For example, the two clusters with the most cells (*Blautia-A*, and *Bifidobacterium*) can be categorized into 10 and 7 sub-clusters, respectively, corresponding to different populations of these genera coexisting in the sample.

### Taxonomic distribution of antibiotic resistance genes

Antibiotics can profoundly impact the gut microbiome in terms of species composition, microbial metabolic activity, and antibiotic-resistant gene (ARG) abundance(Maier et al., 2021). To evaluate ARG distribution among taxa in the fecal microbiome, we searched for ARGs in each cluster (**Fig. 4b**, **Table S12**) by aligning the reads against the Comprehensive Antibiotic Resistance Database (CARD)(Alcock et al., 2019; McArthur et al., 2013). We filtered the alignments by mapping score (Bowtie2 output SAM MAPQ >=42), selected the ARGs for protein coding, and identified 14 ARGs from 15 genus clusters, with mechanisms including antibiotic target alteration, protection, replacement, inactivation, and efflux. ARGs with accession numbers ARO:3004480, ARO:3004601, and ARO:3000190 are most prevalent among the 14 ARGs and were identified in 10, 9, and 6 genus clusters, respectively. Species from *Bifidobacterium*, *Blautia_A*, *Collinsella*, and *Anearobutyricum* carry the most ARGs, at respective counts of nine, eight, six and six. Based on those finding, we predict *Bifidobacterium* potentially has strong resistance (11.2 ARGs read per million reads) to rifampicin and peptide antibiotics, consistent with prior findings(Lokesh et al., 2018). *Dorea* also has high potential resistance to aminoglycoside antibiotic (11.1 ARGs read per million reads) and tetracycline antibiotics (6.7 ARGs read per million reads).

### Functional annotation of gene clusters detected in fecal microbiome genera

Biosynthetic pathways are often encoded as gene clusters that allow cells to acquire the ability to synthesize new molecules(Cimermancic et al., 2014; Wilson et al., 2014). Many gene clusters have already been observed and characterized for function, allowing this information to be annotated to single cell datasets based on detection of key pathway genes, such as MetaCyc (Caspi et al., 2020, 2013; Karp and Caspi, 2011) and KEGG(Kanehisa, 2019, 2000; Kanehisa et al., 2021). Using MetaCyc, in the 95 genera groupings found in our fecal microbiome, we identified 194 gene clusters belonging to 29 classes in 5 super classes, with biosynthetic functions including generation of energy precursors, degradation utilization and assimilation, transport, and macromolecule modification. Additionally, we found that different taxa possess distinct pathways, as might be expected on their unique ecological niches (**Fig. 4c**, **Table S13**). Even within a similar pathway type, different genera have different functions, such as amino acid metabolism. For example, *Blautia_A* possess the pathway to produce arginine, aspartate, ornithine, lysine, methionine, serine, and tryptophan; *Bifidobacterium* to synthesize the branched amino acids isoleucine, serine, and valine; *Akkermansia* to synthesize arginine, isoleucine, valine, and branched amino acid; and *Anaerobutyricum* to synthesize ornithine and methionine. Different genera also utilize distinct carbohydrate sources, with pathways for glucose, galacturonate, lactose, trehalose, sucrose, galactose, stachyose, rhamnose, and mannose all detected in the microbiome. Glucose degradation was identified in *Bifidobacterium*; sucrose degradation was seen in *Agathobacter*, *Anaerostipes*, *Coprococcus*, *Ruminococcus_D* and *Streptococcus*; and, starchyose degradation was detected in *Blautia_A*, *Coprococcus*, *Fusicatenibacter*, *KLE1615*, and *Roseburia*. The ability to unambiguously link functional properties to community members is useful for unraveling the web of pathways that comprise all microbiomes and, ultimately, should aid in the engineering of microbiomes to improve gut health.

### Taxonomic distribution of nutrient biosynthesis pathways

The gut microbiome is the source of vitamins and other nutrients important to health(Krautkramer et al., 2021; Pham et al., 2021). We identified 28 vitamins, cofactors, and carrier biosynthesis pathways in the fecal genome atlas, responsible for producing several vitamins and their precursors, including pantothenate (vitamin B5), adenosylcobalamin (vitamin B12), folate (vitamin B9), riboflavin (vitamin B2), thiamine (vitamin B1), biotin (vitamin B7), pyridoxal 5’-phosphate (active form of vitamin B6), nicotinamide adenine dinucleotide (NAD) and 1,4-dihydroxy-6-naphthoate (precursor of menaquinones or vitamin K2). Riboflavin is produced by 18 genera, including *Agathobaculum*, *Barnesiella*, *Parabacteroides*, and *Coprococcus*. Thiamin pathways exist in 13 genera, including *Bacteroides*, *Bifidobacterium*, *Faecalibacterium*, and *Phocaeicola*, and 19 clusters are detected for folate transformation, including, *Acetatifactor*, *Alistipes*, *Bacteroides*, *Barnesiella*, *Bifidobacterium*, *Blautia_A*, *Eubacterium_F*, *Faecalibacterium*, *Gemmiger*, *Mediterraneibacter*, *Phascolarctobacterium*, *UMGS1375*, and *Phocaeicola*. We also detected 21 clusters containing the pantothenate biosynthesis pathway, including *Acetatifactor*, *Alistipes*, *Bacteroides*, and *Gemmiger*, and that *Alistipes* also synthesizes vitamin K2. These findings show that EASi-seq can characterize nutrient interactions between microbiome members, and between the microbiome and its host.

### Single cell atlas of a coastal sea water microbiome

Environmental microbiomes play important roles in the global ecosystem(Azam and Malfatti, 2007; Sokol et al., 2022), for biogeochemical cycling of elements(Bianchi et al., 2018; Kappler et al., 2021; Klotz et al., 2011; Ramond et al., 2022), metabolism of greenhouse gases(Basiliko et al., 2013; Horn et al., 2006; Sanford et al., 2012), soil fertility(Hayat et al., 2010), and biodegradation(Leahy and Colwell, 1990). Compared to human microbiomes, environmental microbiomes are more diverse and difficult to culture(Hofer, 2018; Pedrós-Alió and Manrubia, 2016). Thus, just as single cell atlases can reveal unique information about human microbiomes, so too can they provide insight into the microbiomes of the environment. To demonstrate the utility of EASi-seq for analyzing environmental microbiomes, we applied it to seawater samples collected from the San Francisco coastline. We isolated the cells via filtration(Lan et al., 2017) (**Fig. S16b**) and processed them with EASi-seq to obtaining 329,470,030 paired-end reads. Quality filtering and further filtration based on classification rate in Kraken2 (**Fig. S23**) yields 3417 cells with an average of 21,062 reads. Using the TDA, we discover 876 genus clusters (**Fig. 5a**, **Table S14-15**), of which 3395 cells are bacteria, and 22 are archaea (**Fig. 5b**). The most abundant bacteria phyla are *Proteobacteria* (2438 cells), *Bacteroidota* (556 cells), *Actinobacteriota* (146 cells), *Verrucomicrobiota* (48 cells), *Firmicutes_A* (34 cells), and *Firmicutes* (22 cells). The archaea include *Thermoproteota* (12 cells), *Halobacteriota* (6 cells), and *Thermoplasmotota* (4 cells). To demonstrate the diversity of the captured community, we constructed a phylogenetic tree using the genus level identification of the cells (**Fig. 5b, center**). Within the 668 identified genera, the top genera by abundance are *Halioglobus* (810 cells), *Sediminibacterium* (218 cells), *Pelagibacter* (190 cells), *Azonexus* (170 cells), *Luminiphilus* (154 cells), and *Amylibacter* (105 cells). This composition is consistent with previous studies of ocean microbiomes(Lan et al., 2017; Rusch et al., 2007; Venter et al., 2004).

**Figure 5.**
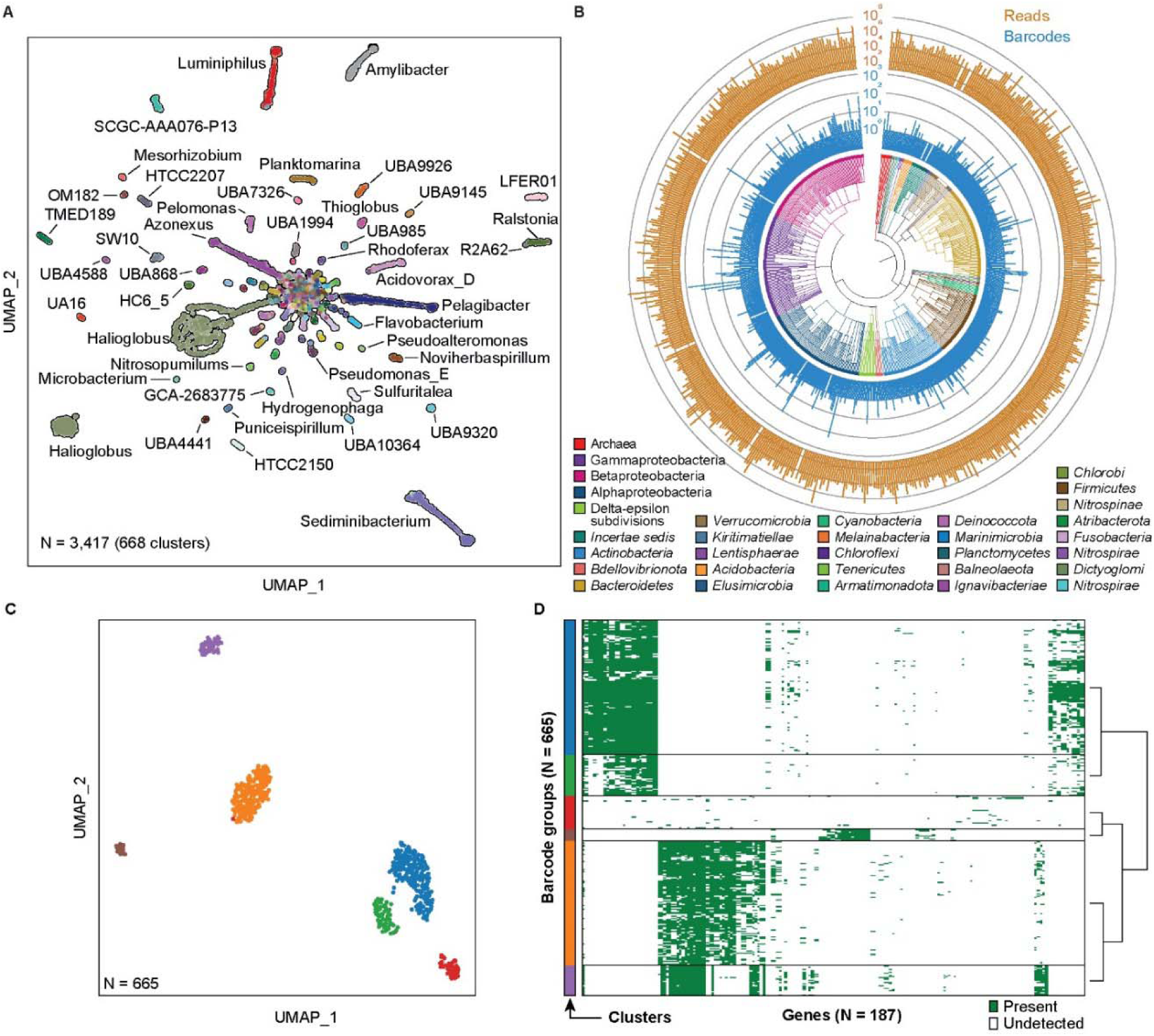
Coastal sea water microbiome genome atlas. (**A**) UMAP based on single cell genera clustering, with points colored according to Kraken2 annotation. (**B**) Phylogenetic analysis of the barcode groups. The phylogenetic tree branches are colored by phylum, except in the case of Archaea (kingdom), Gamma/ Beta/ Alphaproteobacteria (classes all part of the phylum *Pseudomonadota*), and *incertae sedis* (members with ambiguous TDA classification). The inner bar plot (*blue*) shows the barcode count for the corresponding genus, and the outer bar plot (*gold*) shows the total read counts for the corresponding genus. (**C**) UMAP subclustering of the genus *Halioglobus*, from (**A**), which has the highest barcode count. 3873 genes are identified from the 809 barcode groups using HUMANN 3.0 with UniRef90 database. After the genes and barcode groups are filtered (minimum cell counts of a gene = 5 and minimum gene count of a cell = 5), the vector containing 665 barcode groups and 298 genes are used to generate the UMAP. (D) The gene distribution in the *Halioglobus* barcodes. The barcodes are grouped by the cluster in (**C**).

### Single cell gene distribution in *Halioglobus*

To demonstrate the ability to analyze the gene distribution at single cell level, we analyzed the genus cluster with highest abundance, *Halioglobus*, which accounts for 810 barcode groups (23.7% of the 3,417 total counts). The genus *Halioglobus* belongs to the class Gammaproteobacteria and family *Halieaceae*, which is characterized as Gram-negative, non-endospore-forming, aerobic, oligotrophic, and mesophilic bacteria. The family is exclusively isolated from marine environments and is one of the major bacteria groups in coastal or open ocean environments(Han et al., 2019). Although a few isolated *Halioglobus* species isolates have been reported(Han et al., 2019; Kim et al., 2017; Li et al., 2020a, 2020b; Park et al., 2012), the heterogeneity within a *Halioglobus* population has never been studied. Within the single cell cluster, we first annotate genes using HUMAnN3(Beghini et al., 2021). After filtering based on the gene count and cell counts (minimum cell counts per gene = 10 and minimum gene counts per cell = 10), the gene presence/absence matrix of the *Halioglobus* barcode groups are grouped into 6 clusters by Leiden algorithms(Traag et al., 2019) (**Fig. 5c**). The gene presence/absence matrix of each cluster is shown in **Fig. 5d**. This result suggests the *Halioglobus* genus in our sample is a heterogeneous population and indicates that EASi-seq is suitable for the analysis of a heterogeneous population, which could potentially be used for more detailed single-cell-resolution pan-genome analysis.

## Discussion

Microbes play key roles in all ecosystems and are important to human health. While they comprise the most diverse forms of life on the planet(Hug et al., 2016) there are few tools available for sequencing them at single-cell resolution. Additionally, while tools for single mammalian cell genomics have become widespread(Stuart and Satija, 2019), analogous tools for microbes have lagged, due to the technical challenges of isolating and sequencing them in the numbers required to characterize diverse microbiomes. With these realizations in mind, we developed a workflow for efficient microbe sequencing using Mission Bio’s commercial single-cell platform. This instrument is broadly distributed and accessible to non-experts, and therefore constitutes an opportunistic foundation on which to build a single microbe sequencing technology. A major element enabling this has been a simple and general bulk technique to purify single-cell genomes in hydrogels that are compatible with the instrument. Our lysis procedure is applicable to all microbe types, including archaea, bacteria, and fungi, and the commercial microfluidics allow high throughput and efficient single-cell barcoding, to obtain unbiased sequencing for tens of thousands of cells in a sample per run.

The data generated by EASi-seq is unique in that reads are grouped at the level of single cells. By contrast, the dominant method of metagenomic sequencing discards single cell information and captures the sequence data as a mixed pool of short reads. This mixed pool output necessitates complex bioinformatic approaches for strain variation analysis or contig reconstruction that cannot exploit single-cell information. EASi-seq not only overcomes these limitations, but also provides unique methods validating bioinformatic algorithms by comparing results with EASi-seq datasets(Smith et al., 2022). In addition to developing a novel approach for obtaining single-cell data, we also develop novel bioinformatic approaches that exploit the data’s single-cell structure. These include ways to allow cells to be clustered by similarity, aggregation of the reads within a cluster to increase genome coverage, annotate phylogeny and genes, and to scan genomes for genetic elements of interest. By enabling the construction of detailed cell atlases that capture the overall species demographics of a microbiome, EASi-seq affords new opportunities for characterizing the interaction webs inherent to these systems that are near impossible to obtain with metagenomic techniques. With the ability to integrate EASi-seq and metagenomic reads, EASi-seq can provide a complementary viewpoint to metagenomics.

There still remain aspects of the EASi-seq method that can be improved. Importantly, the coverage per barcode is low, which is caused by three reasons. First, before droplet barcoding, the genomic DNA is fragmented by Nextera-like tagmentation, which leads to only 50% of the genomic fragments being viable for barcoding PCR(Picelli et al., 2014). To overcome this inefficiency, we anticipate that future implementations of EASi-seq can increase the complements of adaptors(Tan et al., 2018) or use single-adaptor transposition and uracil-based adapter switching within the barcoding PCR step(Mulqueen et al., 2021). Second, the heterogeneous genome sizes of microbes require different amounts of transposase to achieve the appropriate fragment size(Rodríguez-Gijón et al., 2022). Although we did extensive optimization, one concentration does not fit all needs. For certain genomes, the transposase concentration could either be too high (for smaller genomes) and generate fragments that are below the size-selection threshold or too low (for larger genomes) and produce long fragments incompatible with downstream processing. Third, in adapting the protocol to directly integrate into a commercial device, it was necessary to utilize the barcoding beads from the Tapestri V2 reagent kit. The beads’ barcoding primer has a 15 bp constant region with a melting temperature of 48°C. While we used this sequence as the forward priming site in the barcoding PCR, a higher temperature of 55°C was used as the anneal temperature to avoid random priming. This may lower efficiency in the PCR step. Future optimization can involve development of barcoding beads with improved primers having an elevated melting temperature. Finally, we suspect that coverage can be also improved with an additional single genome amplification step prior to the tagmentation, which can be achieved either in droplet(Zheng et al., 2022) or in hydrogel beads(Nishikawa et al., 2022). Such improved coverage will greatly advance the application of EASi-seq.

Even in its current form, EASi-seq represents a highly accessible platform technology for generating detailed and comprehensive single-cell genome atlases independent of isolation and culturing. Such atlases will have a broad and sustained impact on microbiology, similar to what has been accomplished for mammalian cells. Because we build our workflow on a commercial architecture that is constantly adding features, many of the same improvements and innovations may carry over to microbiomes. For example, after the first demonstrations of mammalian cell DNA and RNA sequencing, multiomic approaches were built on top of the original technologies. These include the ability to measure surface and internal proteins, characterize epigenetic signatures and genome structure, and integrate spatial data(Ogbeide et al., 2022). For example, microbial RNA-seq is possible using universal cDNA methods amenable to single cell barcoding and would thus allow addition of transcriptional state measurements with EASi-seq. Using oligonucleotide-labeled binders like including antibodies, lectins, and aptamers, microbes can be stained prior to EASi-seq, allowing for recording proteomic and serotype signatures in a manner similar to Ab-seq(Shahi et al., 2017), DAb-seq(Demaree et al., 2021), CITE-seq(Stoeckius et al., 2017), inCITE-seq(Chung et al., 2021), and INS-seq(Katzenelenbogen et al., 2020). Similarly, the lysis and molecular biology processes of EASi-seq should carry over to DNA viruses and, with the implementation of reverse transcription, RNA viruses, holding potential for single virus genome atlasing.

## Materials and Methods

1. Microbiome samples processing
a. Synthetic community ZymoBIOMICS standard (Zymo, D6300) was stored at-80°C until use. 100 µL of ZymoBIOMICS was washed with 4 mL of PBS for 3 times to remove the storage buffer. The cell density is measured with Countess™ cell counting slides (Thermo Fisher, C10228) using an EVOS microscope. After counting, cells were resuspended to a final concentration of 100 million per mL in PBS. All twenty-two *E. Lenta* strains (**Table S3** list of *E. lenta* strains) were cultured in appropriate media(Bisanz et al., 2020)and equally mixed based on CFU counting in culture media. The cell mixture is stored at-80°C until use. Before processing, thawed cells were washed 3 times to remove the storage media and filtered with 5 µm syringe filter to remove cell aggregates. After cell counting, the cells were resuspended to 100 million per mL in PBS.
b. Human microbiome and cell isolation Fecal sample from health donor is stored at-80=°C until use. Cell isolation was performed according to previously reported protocol(Hevia et al., 2015).
c. Ocean water microbiome and cell isolation Sea water was collected at Pacific coastline near San Francisco (GPS coordinate: 37.7354373 N, 122.5081862 W) by submerging a 1000 mL sterile bottle into the ocean. The sea water was transferred to the lab on ice. The cells were enriched according to the published protocol(Lan et al., 2017). Briefly, the sea water was first filtered through a 50 µm cell strainer (Corning, 431752) to remove sands or other large particles. The suspension was then filtered by a 0.45 μm vacuum filter (Millipore, SCHVU01RE) to capture the cells on the membrane. The membrane was cut off from the filter with a sterile razor blade and transferred a 15 mL centrifuge tube with 5 mL PBS. The cells were released from the membrane by vortexing the tube at maximum speed for 2 min. The cells were washed with 10 mL PBS for 3 times and passed through a 5 μm syringe filter to remove remaining virus or large particles. The cells were resuspended to 100 million per mL in PBS.
2. Microfluidics device fabrication Microfluidics devices were fabricated with standard photolithography and soft lithography method. Custom device fabrication is not necessary for the single cell sequencing using Mission Bio Tapestri but used for workflow optimization. Master photomask was designed using AutoCAD and printed at 12,000 DPI (CAD/Art Services, Bandon, OR). To make the master structure, SU8 Photoresist (MicroChem, SU8 3025 and SU8 3050) were spin coated on three-inch silicon wafers (University Wafer), soft baking at 95°C for 10 to 20 min, UV-treated through the photomasks for 3 min, hard baked at 95°C for 5 to 10 min and developed in propylene glycol monomethyl ether acetate (Sigma Aldrich). For the microfluidic devices, poly(dimethylsiloxane) (PDMS) (Dow Corning, Sylgard 184) and curing agent were mixed in 10:1 ratio, degassed and poured over the master structure, baked at 65 °C for 4 h to cure, and peeled off from the wafer. After hole punched with a 0.75 mm biopsy puncher, the devices were plasma treated and bonded to glass slides. The channels were treated with Aquapel (PPG industry) to for hydrophobic surface and dried by baking at 65°C for 10 min.
3. Single cell genomic DNA isolation in hydrogel beads
a. Cell encapsulation in hydrogel beads 500 µL cell suspension (100 million per mL in PBS) was mixed with 500 µL hydrogel precursor solution (12% acrylamide, 1% BAC, 20 mM Tris, 0.6% sodium persulfate, and 20 mM NaCl in H_2_O) in a 15 mL centrifuge tube. 1 mL HFE 7500 with 2% surfactant (008-FluoroSurfactant, RanBiotechnologies) was added to the cell/hydrogel precursor mixture. Emulsion was formed by passing the oil/aqueous mixture 5 times through a 20 gauge syringe needle. 20 µL of TMEDA (tetramethylethylenediamine, Sigma) was added into the emulsion and the emulsion was incubated at 70 °C for 30 min and at room temperature for overnight for gelation. The emulsion can be stored at 4°C for up to 1 week. The emulsion was centrifuged at 1000 RCF for 1 min and the bottom oil layer was removed by using a gel loading tip. 1 mL of 20% PFO (1H,1H,2H,2H-perfluoro-1-octanol, Sigma, 370533) and 5 mL of PBST buffer (0.4% tween 20 in PBS) were added into the emulsion. The mixture was vortexed at maximum speed for 1 min to break the emulsion and centrifuged at 1000 RCF for 5 min. Any remaining oil was removed by pipetting through a gel-loading tip.
b. Hydrogel size selection Differential velocity centrifugation was performed to select the hydrogel beads from previous step within the diameter between 5 to 15 µm. 500 µL hydrogel beads were resuspended in 14 mL high density buffer (40% sucrose in PBS with 0.4% tween 20). First, the beads were centrifuged at 1000 RCF for 5 min to pellet large gels. The supernatant was transferred to a new 15 mL tube and centrifuged at 3000 RCF for 10 min to pellet the right sized beads. The supernatant (still containing beads smaller than 5 µm) was discarded and the pelleted beads were washed 3 times with PBST to remove the high-density buffer. Typically about 300 µL of size selected beads can be recovered.
c. Cell lysis in hydrogel beads 100 µL of size selected beads were treated in 1 mL cell wall digestion buffer (TE buffer solution containing 2.5 mM EDTA, 10mM NaCl, 2U zymolyase, 5 U Lysostaphin, 50 U mutanolysin, and 20 mg Lysozyme) at 37 °C overnight. The beads were then pelleted by centrifugated at 3000 RCF for 10 min and washed with PBST for 3 times. The beads were then treated in 1 mL protein digestion solution (TE buffer with 4U of Proteinase K, 1% triton X100 and 100 mM of NaCl) at 55 °C for 30 min. Following lysis, the beads were thoroughly washed with PBST, 100% EtOH, and PBST 3 times to ensure complete removal of proteinase K and other chemicals which may inhibit the downstream reactions. The beads were then filtered with 10 µm cell strainer and ready for droplet tagmentation.
4. Single cell tagmentation and barcoding in droplet microfluidics Microfluidic droplet encapsulation, tagmentation, and barcoding PCR were performed on commercial single-cell DNA genotyping platform (Mission Bio, Tapestri) or custom build microfluidic devices with the same functions.
a. Tagmentation reagents 25 µL Tn5-Fwd-oligo GTA CTC GCA GTA GTC AGA TGT GTA TAA GAG ACA G (100 nM, IDT), 25 µL, Tn5-Rev-oligo TAC CCT TCC AAT TTA ACC CTC CAA GAT GTG TAT AAG AGA CAG (100 nM, IDT), and 25 µL Blocked ME Complement /5Phos/C*T* G*T*C* T*C*T* T*A*T* A*C*A*/3ddC/ (200 nM, IDT) and 25 µLTris buffer were mixed well in a PCR tube by pipetting. The mixture was incubated on a PCR thermal cycler with the following program: 85°C for 2 min, cools to 20 °C with a ramping rate at 0.1 °C/s, 20 °C for 1 min, then hold at 4 °C with lid at 105°C. 100 uL of glycerol was added into the annealed oligo. Unloaded Tn5 protein (1 mg/mL, expressed by QB3 MacroLab, Berkeley, CA), dilution buffer (50% Glycerol, 100 mM NaCl, 0.1 mM EDTA, 1 mM DTT, and 0.1% NP40 in 50 mM Tris-HCl pH 7.5 buffer), and the pre-annealed adapter/glycerol mix were mixed at 1:1:2 ratio by pipetting. The mixture was incubated at room temperature for 30 min then stored at-20 °C until use. For droplet tagmentation, equal amount of assembled Tn5 and tagmention buffer (10 mM MgCl2, 10 mM DTT in 20 mM TAPS pH 7.0 buffer) were mixed.
b. Droplet tagmentation In the first droplet step, the tagmentation reagents (0.125 mg/mL assembled Tn5, 10 mM MgCl2, and 10 mM DTT in 20 mM TAPS pH 7.0 buffer) and the genomic DNA in hydrogel beads (equivalent to 3 million cells per mL) in 10 mM MgCl_2_, 1% NP40, 17% Optiprep, and 20 mM TAPS pH 7.0 buffer were co-flowed in the microfluidic devices to form droplets. In case of using Tapestri, the MissionBio Tapestri DNA cartridge and a 0.2 mL PCR tube were mounted onto the Tapestri instrument. 50 µL beads solution, 50 µL tagmentation reagents, and 200 µL encapsulation oil were load in the cell well (reservoir 1), lysis buffer well (reservoir 2), and encapsulation well (reservoir 3) on the Tapestri DNA cartridge, respectively. The Encapsulation program was used for droplet generation. The droplets were collected into a PCR tube. For custom build microfluidic device, the beads solution, the tagmentation reagents, and 5% (w/w) PEG-PFPE surfactant (Ran Biotechnologies) in HFE 7500(3M) were loaded into three syringes and placed on syringe pumps. The syringes were connected to the co-flow droplet generator device via PTFE tubing. The pumps were controlled by a Python script (https://github.com/AbateLab/Pump-Control-Program) to pump bead solution at 200 µL/h, tagmentation reagents at 200 µL/h and oil at 600 µL/h to generate droplets. The droplets were collected into PCR tubes. The droplets generated by either method are incubated at 37°C for 1 h, 50°C for 1h, and hold at 4°C to ensure hydrogel melting and Tn5 complete reacting.
c. Droplet barcoding PCR The tagmentation droplets from the previous were merged with PCR reagents and barcode beads for barcoding with either Tapestri or custom build microfluidic devices. In case of using Tapestri, 8 PCR tubes and DNA cartridge were mounted onto the Tapestri instrument. Electrode solutions were loaded into electrode wells (200 µL and 500 µL in reservoirs 4 and 5, respectively). After running the Priming program, 5 µL of reverse primer (GTC TCG TGG GCT CGG AGA TGT GTA TAA GAG ACA GTA CCC TTC CAA TTT AAC CCT CCA, 100 µM, IDT) was mixed with 295 µL Mission Bio Barcoding Mix and loaded into PCR reagent well (reservoir 8) of the DNA cartridge. The droplets from previous step (∼80 µL), 200 µL of V2 barcoding beads, and 1.25 mL of Barcoding oil were loaded into cell lysate well (reservoir 6), barcode bead well (reservoir 7) and barcode oil well (reservoir 9), respectively. The droplets were merged with barcoding beads and PCR reagents by the Cell Barcoding program. The resulting droplets were collected into the 8 PCR tubes. In case of using custom build microfluidics, the device was first primed by filling electrode solution (2M NaCl solution) into the electrode and the moat channels. 500 µL PCR reagents containing 1.67X Q5® High-Fidelity Master Mix (NEB, M0515), 0.625 mg/mL BSA, 1.2 µM reverse primer (GTC TCG TGG GCT CGG AGA TGT GTA TAA GAG ACA GTA CCC TTC CAA TTT AAC CCT CCA) were loaded into a 1 mL syringe. 200 µL Mission Bio V2 barcoding beads washed with Tris buffer (pH 8.0) and resuspended in 10 mM Tris buffer containing 3.75% tween 20, 2.5 mM MgCl2, 0.625 mg/mL BSA. The beads were centrifuged at 1000 RCF for 1 min and the supernatant was removed. The remaining bead slurry (∼110 uL) was loaded into PTFE tubing connected to a 1 mL syringe filled with HFE 7500 oil. The droplets after tagmentation were loaded into a 1 mL syringe. The three syringes and two syringes filled with 10 mL of 5% (w/w) PEG-PFPE surfactant (Ran Biotechnologies) in HFE 7500(3M) HFE 7500 were connected to the microfluidic devices. The follow rates are as follows: tagmenation droplets 55 µL/h, spacer oil 200 µL/h, PCR reagent 280 µL/h, barcode beads 148 µL/h, and droplet generation oil 2000 µL/h. To merge the tagmentation droplet, the electrode near the droplet generation zone was charged with an alternating current (AC) voltage (3 V, 58kHz). And the moat channel was grounded to prevent unintended droplet coalescence at other locations on the device. The merged droplets were collected into PCR tubes. The droplets collected in the merging step were treated with UV for 8 min (Analytik Jena Blak-Ray XX-15L UV light source) and the bottom layer of oil in each tube were removed using a gel loading tip to leave up to 100 µL of droplets. The tubes were placed on PCR instrument and thermo-cycled with the following program: 10 min at 72°C for 1 cycle, 3 min at 95°C for 1 cycle, (15 s at 95°C, 15 s for 55°C, and 2 min at 72°C) for 20 cycles, and 5 min at 72°C for 1 cycle with the lid set at 105°C.
d. Barcoded Amplicon purification The thermal cycled droplets in the PCR tubes were carefully transferred into two 1.5 mL centrifuge tubes (equal amount in each). If there were visible merged large droplets present, they were carefully removed using a 2 µL pipette. 20 µL PFO were added into each tube and mixed well by vortex. After centrifuging at 1000 RCF for 1 min, the top aqueous layers in each tube were transferred into new 1.5 mL tubes without disturbing the bead pellets and water was added to bring the total volume to 400 µL. The barcoding product was purified using 0.7X Ampure XP beads (Beckman Coulter, A63882) and eluted into 50 µL H2O and stored at-20°C until next step. The concentrations of the barcoding product were measured with Qubit™ 1X dsDNA Assay Kits (ThermoFisher, Q33230).
5. Barcoding sequencing library preparation and sequencing
a. Library prep and QC The sequencing library were then prepared by attaching P5 and P7 sequences to the barcoding products using Nextera primers (**Table S2**). The library PCR reagents containing 25 uL Kapa HiFi Master mix 2X, 5 uL Library P5 index primer (4 uM), 5 uL Library P7 index primer (4 uM), 10 uL purified barcoding products (normalized to 0.2 ng/uL), and 5 uL of nuclease free water were thermal cycled with the following program: 3 min at 95°C for 1 cycle, (20 s at 98°C, 20 s for 62°C, and 45 s at 72°C) for 12 cycles, and 2 min at 72°C for 1 cycle. The sequencing library was purified with 0.69X Ampure XP beads and eluted into 12 uL nuclease-free water. The library was quantified with Qubit™ 1X dsDNA Assay Kits and DNA HS chips on bioanalyzer or D5000 ScreenTape (Agilent, 5067-5588) on Tapestation (Agilent, G2964AA). The libraries were pooled and 300 cycle pair-end sequenced by Illumina MiSeq, NextSeq or NovaSeq platform.
6. Sequencing file barcode extraction and single cell read file preparation Raw sequencing FASTQ files were processed using a custom python script (mb_barcode_and_trim.py) available on GitHub (https://github.com/AbateLab/MissonBioTools) for barcode correction and extraction, adaptor trimming, and grouping by barcodes. For all reads, combinatorial cell barcodes were parsed from Read 1, using Cutadapt (v2.4)(Martin, 2011) and matched to a barcode whitelist. Barcode sequences within a Hamming distance of 1 from a whitelist barcode were corrected. Reads with valid barcodes were trimmed with Cutadapt to remove 5′ and 3′ adapter sequences and demultiplexed into individual single-cell FASTQ files by barcode sequences using the script demuxbyname.sh from the BBMap package (v.38.57)(Chaisson and Tesler, 2012).
7. Reference based single cell data analysis
a. ZymoBIOMICS Microbial Community Standards The reference genome FASTA files of the ten species of Zymo BIOMICS Microbial Community Standards provided by Zymo Research Corporation (https://s3.amazonaws.com/zymo-files/BioPool/ZymoBIOMICS.STD.refseq.v2.zip). The FASTA files were combined and Bowte2 index were built using Bowtie2-build command. The reads in single-cell FASTQ files were aligned to reference genomes using Bowtie2 (v 2.3.5.1) with default setting(Langmead and Salzberg, 2012). The overall alignment rates for each barcode were collected from the log files. The barcode groups less than 50% overall coverage rate were removed. Each barcode group’s coverages, numbers of mapped reads, covered bases, and mean depths of 10 corresponding species were calculated using Samtools v1.12 (samtools coverage) with default setting(Li et al., 2009). The purity of each barcode group was calculated as the percentage of reads that aligned to a dominant species. For the rarefaction analysis, 10,000 reads were randomly sampled from the SAM file of each barcode group. The coverage was calculated after each read sampling using Samtools.
b. Strain abundance estimation for synthetic community with 22 *E. lenta* strains The reference genomes of the 22 *E. lenta* strains were downloaded from NCBI (**Tabel S3**). The reads in single-cell FASTQ files were aligned to reference genomes using Bowtie2 (v 2.3.5.1) (Langmead and Salzberg, 2012)with-a setting to report all matches. The overall alignment rates for each barcode were collected from the log files. The barcode groups with less than 50% overall coverage rate were removed. The probabilities of each alignment were calculated with parseAlignment command from BitSeq (v 1.16.0)(Glaus et al., 2012)
8. Taxonomic Discovery Algorithm
a. TDA validation using simulation data 100 species were randomly selected from the NCBI assembly metadata file (ftp://ftp.ncbi.nlm.nih.gov/genomes/genbank/bacteria/assembly_summary.txt). The reference genome FASTA files were downloaded using the corresponding link in the metadata file (**Table S4**). Simulated pair-end read files were generated using a Python script according to the following rules. 1. 100 barcode groups were generated for each species. 2. The reads are 150 bp paired end. 3. The amplicon length is in the range of 400-1000 bp. 4. Each barcode group has 0-49% percent of contamination reads. 5. The contamination reads were generated from the other 99 species. 6. Each barcode has 1,000-10,000 pair-end reads. 3 taxonomic classifiers were chosen for evaluation: Kraken2/Bracken v (Lu et al., 2017; Wood et al., 2019) with PlusPF database(https://benlangmead.github.io/aws-indexes/k2, Version: 1/27/2021), Kaiju v(Menzel et al., 2016) with its standard database and MetaPhlAn v(Beghini et al., 2021) with its standard database. All the pair-ended barcode group FASTA files were profiled using the three classifiers. The results were grouped and analyzed in Python. The predicted taxa purity was the abundance of the dominant taxa in each barcode group. The barcode filtering based on purity was performed using thresholds ranging from 50% to 99% purities. The average after-filtering purity was the mean purity of all the barcodes that passed a certain threshold and after-filtering barcode counts was the barcode count of which passed a certain threshold. The UMAP clustering was performed with the genus abundances of all the barcode groups. The identity of each cluster was assigned with the most abundant taxa. The identification accuracy was calculated as the percentage of barcodes with the correct genus identification.
b. TDA analysis of single cell sequence data The single cell sequencing barcode group FASTQ files of ZymoBIOMICS, Human microbiome and the sea water microbiome samples were analyzed using TDA with Kraken2/Bracken as the taxonomic identifier. For the Zymo BIOMICS sample, Kraken2 PlusPF database (https://benlangmead.github.io/aws-indexes/k2, Version: 1/27/2021) was used, while for human microbiome and sea water microbiome, Kraken2 GTDB database (https://gtdb.ecogenomic.org/tools, Release 95) was used. The reads in each barcode group were first classified by Kraken2, and the abundances at genus and species level were re-estimated with Bracken using default threshold setting. The percentages of the mapped reads were extracted from the Kraken2 output files of barcode groups. The purities were calculated as the abundance of the dominant genus in the barcode groups. The data was filtered according to percentage of mapped reads and genus-level purity. The taxa abundance profiles of the remaining barcodes were combined and UMAP clustering was performed using The Scanpy toolkits(Wolf et al., 2018) in Python script. The taxa of each barcode group were assigned to the most abundant one.
9. Metagenomic sequencing and assembly
a. ZymoBIOMICS community The metagenomic sequencing data of ZymoBIOMICS Microbial Community Standards D6300 (batch ZRC195925) was provided by Zymo Research Corporation. The reads were assembled using SPAdes-3.15.3 with ‘--meta’ setting(Prjibelski et al., 2020).
b. Human microbiome The human fecal sample was collected from a healthy adult donor under a UCSF IRB approved protocol (#14-13821). The sample was deposited into a commode specimen collection system and aliquoted into 2mL cryovials with DNA/RNA shield (Zymo). For bulk metagenomic sequencing, the sample was extracted using the ZymoBiomics 96 MagBead DNA kit. The sequencing library was prepared using the Nextera XT protocol and sequenced using an Illumina Nova-Seq with 2×140 chemistry at the Chan Zuckerberg Biohub. Metagenomic reads were quality-filtered using FastP (v. 0.20.0)(Chen et al., 2018).
10. Comparison between metagenomic and single cell sequencing. The genus abundances of the human microbiome metagenomic data and the pooled single cell sequence file were analyzed using Kraken2 and Bracken. The results were plotted as a scatter plot with triangle markers. For any genus with a barcode group associated, a round marker of the genus was added and its size is proportional to the barcode counts.
11. Single cell sequencing data integration with metagenomics To integrate the metagenomic dataset, the contigs assembled from metagenomic sequencing (ZymoBIOMICS and human microbiome sample) were treated as individual barcodes and processed with TAD. The metagenomic reads were first aligned to the assembled contigs using Bowtie2 v2.3.5.1(Langmead and Salzberg, 2012).
12. Clustered barcode groups analysis
a. Cluster assembly and evaluation Single cell barcodes of UMAP clusters were combined using concatenate command (cat) in the Linux system into single FASTQ files. The pair-end reads associated with barcodes that belong to the same UMAP clusters were grouped by Seqtk toolkit (https://github.com/lh3/seqtk) (seqtk subseq) into single FASTQ files. The assemblies were conducted with all reads associated to both single cell sequencing and metagenomic contigs of each UMAP cluster using Spades v 3.15.3(Prjibelski et al., 2020) with ‘--careful’ setting. The assembled contigs were evaluated using Quast v 5.0.2 with or without reference genome input. To calculate the clustering error rate, all the reads associated to a cluster were mapped to the corresponding reference genome, the percentage of the reads that were not aligned was considered as the error rate. Pathway analyses of each cluster was conducted using HUMAnN v 3.0(Beghini et al., 2021) with the default MetaCyc(Caspi et al., 2020, 2014; Karp and Caspi, 2011) database. The pathway abundance files of each cluster were combined and plotted as a heatmap using the Seaborn module in Python. The sub-categorizing of barcode groups in a UMAP cluster was using species abundance estimation. The 2 clusters with the most barcode groups in the human microbiome samples (*Blautia_A*, and *Bifidobacterium*) were further divided into sub clusters by UMAP aggregation with the Kraken2 species abundance estimation.
b. Gene association analysis Comprehensive Antibiotic Resistance Database (CARD) (v 3.1.4)(Alcock et al., 2019) (https://card.mcmaster.ca/download) was downloaded and bowtie2 references were built with botie2-build command(Langmead and Salzberg, 2012). The combined reads associated with each UMAP cluster identified in the human gut microbiome were mapped to the CARD databases using Bowtie2 (v2.3.5.1)(Langmead and Salzberg, 2012). The mapping reads are filtered for MAPQ >= 42 to select the reads without mismatches using SAMTools (samtools view-bS-q 42). After duplicate reads were removed using SAMTools (samtools rmdup-S), the references sequence name (RNAME) of each alignment were extracted from the bam files. The unique genes associated with each UMAP cluster, and their frequencies were generated from the RNAMEs. The relative abundance antibiotic resistance gene is calculated as the unique ARO read count per million total read count. The resistance mechanism associated with antibiotic resistance ontology (AROs) were downloaded from the Comprehensive Antibiotic Resistance Database.
13. Data and code access All sequencing data is accessible at the NCBI Sequence Read Archive (Accession numbers: SUB12874540). Python Jupyter notebooks code used in this paper can be accessed at Abate lab GitHub: (https://github.com/xiangpenglee/EASi-seq.git). The supplementary tables can be accessed at (https://github.com/xiangpenglee/EASi-seq/tree/main/Supplimentary_Data).

## Acknowledgments

The authors are grateful for helpful discussions with Drs. Katie Pollard, Byron Smith, Xiaofan Jin, Jason Shi at Gladstone Institute; Susan Lynch, Xiaoyuan Zhou, Cyrille Delley, Leqian Liu, and Peng Xu at UCSF; Adam Arkin and Fangchao Song at University of California, Berkeley; and Michael Fischbach and Bryan Yu at Stanford University.

## Author Contributions

X.L. and A.R.A. designed the research. X.L., L.X., B.D., and D.W. performed the single cell experiments, X.L. and B.D. analyzed the single cell data, C.N., J.E.B, and P.T.J. provided microbiome samples, X.L., C.N. and J.E.B performed metagenomic experiments and assembly. C.M. provided feedback regarding experimental design and interpretation of data, in addition to help planning of the manuscript. X.L. wrote the initial draft of the manuscript, A.R.A., C. M., C.N., J.E.B, and P.T.J revised the manuscript. All authors read, reviewed, and approved the manuscript.

## Classification

Biological Sciences: Genetics and Microbiology

