## Supplemental Info for "Main Manuscript for Microbiome single cell atlases generated with a benchtop instrument"

**This PDF file includes:**

Discussion S1-S3

Figs. S1 to S23

Table S1

Table S2-S21 description

References (1 to 10)

**Other supporting materials for this manuscript include the following:**

Table S2 to S21

The tables can be accessed at (<https://github.com/xiangpenglee/EASi-seq/tree/main/Supplimentary_Data>)

Supporting Information Text

**Discussion S1. Taxonomic classifier selection for Taxonomic discovery algorithm using simulated single cell sequencing data.**

A key challenge in single microbe sequencing data analysis is that only a fraction of potential microbial species has a reference genome available. Thus, it is often not possible to use read mapping against a genome database to identify a barcode’s taxa. Further, because the average coverage of each barcode group is less than 1%, the likelihood of two different barcodes of the same cell type sharing a common set of reads is low. These factors make it difficult to use a read-based method for clustering barcode groups belonging to the same type of cell. Initial tests with such a method failed to cluster cells with common reads or SNPs between barcode groups.

To identify each barcode group without using reference genomes, we propose the Taxonomic Discovery Algorithm (TDA). In the TDA workflow, each barcode group is treated as a metagenomic sample and its taxonomic abundance is estimated using available taxonomic classifier approaches. Our hypothesis was that different barcode groups belonging to the same cell should be classified to the same taxa by a taxonomic classifier even if each barcode group possess completely different sets of reads. In particular, an ideal taxonomic classifier suitable for TDA should accurately identify each barcode group using a limited number of reads. The taxonomic classifier should also provide accurate abundance of each clade in the circumstances when barcode groups are from droplets that were loaded with more than one cell, so as to enable filtering. There are 3 types of classifier approaches: k-mer based, protein-based, and marker gene-based. These approaches assign taxonomic labels to metagenomic DNA sequences through respectively detecting clade-specific k-mer, protein sequence, or marker gene signatures. For identifying the best approach for TDA use, we evaluated a recent tool from each method: Kraken2(Wood et al., 2019) in pair with Bracken(Lu et al., 2017) (k-mer based), MetaPhlAn(Beghini et al., 2021) (marker gene-based), and Kaiju(Menzel et al., 2016) (protein-based).

Testing used a synthetic sequencing output constructed from the reference genomes of one hundred randomly chosen species (**Table S17**). Every genome was used to generate a hundred barcode groups, each containing 1,000 to 10,000 pair-end reads of 150 bp in length. Simulated barcode groups could also contain up to 49% in contaminating reads (generated from the other ninty-nine species) (**Fig. S7a-b**, **Table S18**).

The generated barcode groups were classified with Kraken2/Bracken, Kaiju, and MetaPhlAn to generate the taxonomic abundance profiles (**Fig. S7c-e**, **Tables S19-21**). We evaluated the correlation between the predicted purity and the known purity of each barcode group (**Fig. S7c-e**) using the Pearson correlation coefficient (r). Kraken2/Bracken showed the highest correlation (r = 0.597), Kaiju showed medium correlation (r = 0.2895) and MetaPhlAn showed the lowest correlation (r = 0.0452). In a low coverage setting (about 1000 to 100,000 reads per barcode group), the chance to find the clade-specific k-mer, protein sequence, and marker genes are proportional to the size of the database. Because there are only approximately one million clade-specific marker genes in the MetaPhlAn database, in most barcode groups, only one or zero marker genes were found. If one marker gene was identified in a barcode group the purity was calculated to be 100%, while barcodes with no identified marker genes had 0% purity. This caused MetaPhlAn predicted purity to show a binomial distribution. The Kaiju database has slightly larger (approximately seventy-seven million) clade-specific protein sequences. Although Kaiju has higher accuracy in predicting barcode group purity than MetaPhlAn, both results are still inferior to Kraken2, which can draw on over ten billion clade-specific k-mers in its database (PlusPF Version: 1/27/2021, <https://benlangmead.github.io/aws-indexes/k2>).

We then mimicked the process of using the predicted purities to filter the barcode groups (**Fig. S7f-g**). With various filtering thresholds, Kaiju showed the highest average after-filtering purity, but the after-filtering barcode counts were significantly decreased. Kraken2/Bracken retained most of the after-filtering barcodes but has lower average after-filtering purity than Kaiju. MetaPhlAn predicted purity showed no filtering effect.

We also generated UMAP clustering(Becht et al., 2019) plots based on the predicted genus abundance in each barcode group (**Fig S7c-e** lower panels). With Kraken2/Bracken and Kaiju, more than 92% and 93% of barcode groups were assigned to the correct genera, while the genus-level identification accuracy of MetaPhlAn was only 77% (**Fig S7h**).

Overall, k-mer based Kraken2/Bracken showed the best performance and was used as the taxonomic classifier in TDA.

Discussion S2. Cell aggregation interaction in frozen human fecal sample

The purity distribution of the human fecal microbiome sample showed three peaks. The peak centered around 100% belongs to pure single cell barcode groups, while the beaks centered at 40% and 55% are for barcode groups containing, respectively, three and two cells. We filtered the barcode groups with >80% purities for single cell analysis (**Fig. S17a**, *blue region*, see discussion in main text). The 1192 barcode groups with purities ranging between 50% to 60% were used to analyze the cell-cell interaction (**Fig. S17a**, *orange region*). We found *Blautia_A* is often associated with *Collinsella* (1000 occurrences), *Bifidobacterium* (50 occurrences), and *Senegalimassilia* (50 occurrences). Additionally, we found that *Collinsella* is found together with *Streptococcus* (12 occurrences) and *Faecalibacterium* (9 occurrences) (**Fig. S17b**).

This cell mixing can occur at various steps in the EASi-seq workflow as artifacts. We addressed this possibility at each step of the process. First, in the cell isolation step, we used density centrifugation to isolate cells away from non-specific cell aggregates or clumping material using a 5 µm filter. This filter is small enough to capture particulate matter, while being large enough to allow small cell groups (such as those with two or three members) to pass through. Second, during the hydrogel encapsulation step, cell overloading can cause multiple cells to be captured in each hydrogel bead. For processing the filtered fecal sample, we used a loading concentration that had consistently showed limited mixing when used for other kinds of samples. Third, in the droplet tagmentation step, cells containing hydrogels are Poisson loaded into droplets. While there is a natural frequency of droplets contain more than one hydrogel by chance, our final densities limited the contribution of this effect. Specifically, at a cell hydrogel density of ~3 million per mL and a droplet size of ~42 µm (~0.04 nL volume), thePoisson distribution predicts a loading rate where 88.25% droplets are empty, 11.03% droplets contain a single hydrogel, and the remaining 0.7% of the droplet population accounts for all doubly loaded and (rare) higher-order loadings. Fourth, in the second droplet step, hydrogel containing droplets are paired and merged with a droplet containing the barcoding bead and PCR reagent. Rarely, two hydrogel droplets can merge with barcoding bead droplets, leading to cell mixing. However, because most of the droplets contain zero cells, mixing events from this mechanism are reason extremely rare. Finally, during the PCR step, droplet merging might also happen during the thermocycling. We ruled out this reason because of the extra surfactant added to stabilize the droplets during PCR and the lack of any significant merging observed on post-PCR inspection of the emulsion.

Thus, given the human fecal sample was processed in the same manner as other biological samples that showed no significant cell mixing, we believe the significant mixing reflects a true cell-cell association phenotype. According to the spatial analysis of the mouse intestinal tract by Metagenomic Plot Sampling by sequencing (MaPS-seq)(Sheth et al., 2019), family *Lachnospiraceae* bacteria (to which *Blautia_A* belongs) showed significant interaction with several other microbial families. This suggests that *Lachnospiraceae* might have properties that allow physical association with other bacteria. Further studies are needed to confirm the suggested cell-cell interactions.

Discussion S3. Human fecal sample EASi-seq identified less genera than metagenomics

Compared to metagenomics, several genera are less represented in EASi-seq. These include *Bacteroides*, *Phocaeicola*, *Parabacteroides*, *Akkermansia*, *and Alistipes* (**Fig. S19**). We suspect this is caused by sample storage and cell isolation process. During the centrifugation-based cell isolation process(Hevia et al., 2015), we use a 5 µm size filter to remove large particles and cellular aggregates. Cells that either stick to fecal particles or have a propensity to form large aggregates might be removed from the sample. This is supported by the observation that certain species of *Bacteroides* can adhere to partially digested food particles(Bäckhed et al., 2005). This bias could also be introduced by the sample storage process at -80 °C, which is known to increase the composition of *Blautia* and *Collinsella*, and decrease that of *Bacteroides* and *Alistipes*(Watson et al., 2019). We can improve the relative abundance accuracy by using fresh collected samples(Zheng et al., 2022).

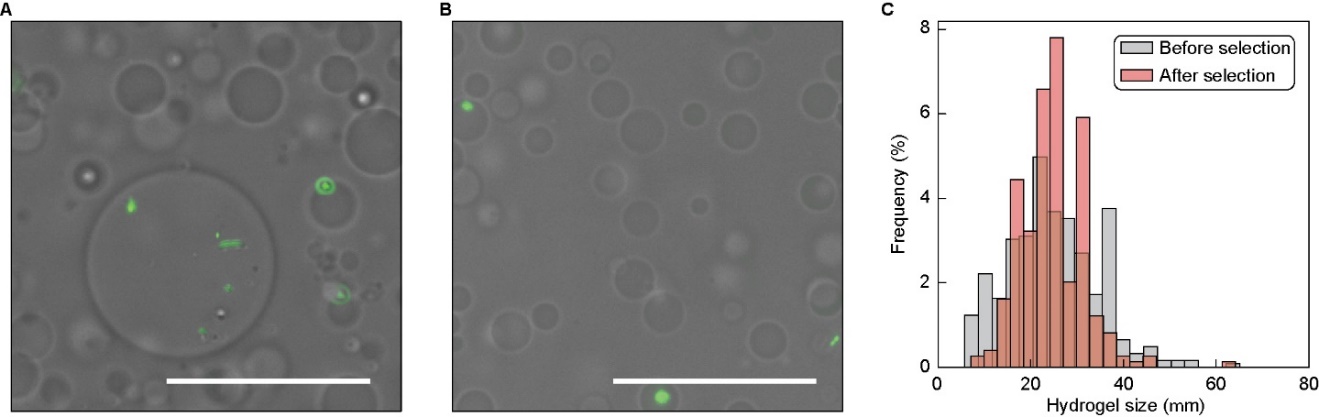

**Fig. S1. Hydrogel beads before and after size selection**.

(**A**,**B**) Images of the hydrogel beads before (**A**) and after (**B**) size selection. The cells embedded within the hydrogel are stained with a DNA stain (SYBR Green I). Scale bar = 50 µm. (**C**) Size distributions of the hydrogel beads before and after size selection.

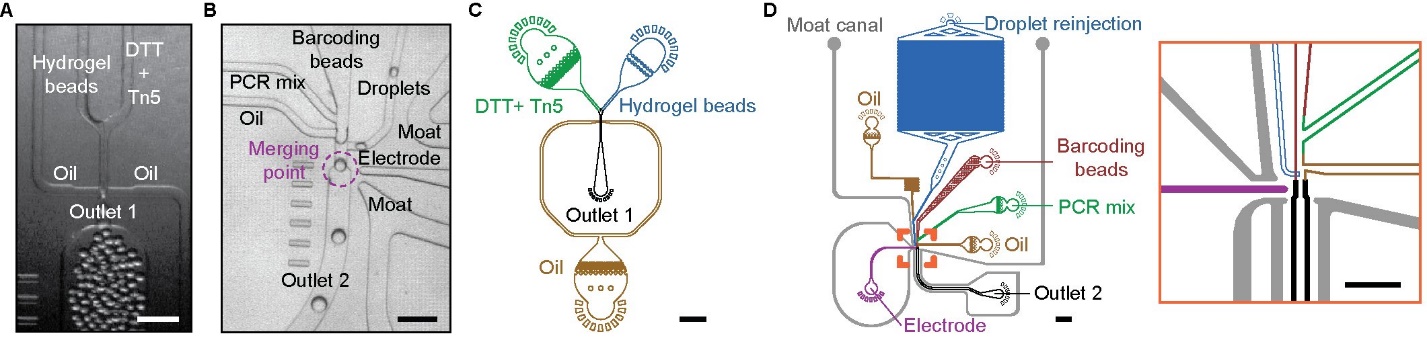

**Fig. S2. Microfluidic device images and schematics.**

(**A**) Image of droplet generation for tagmenation using the Mission Bio Tapestri cartridge. Scale bar = 200 µm. (**B**) Image of tagmented droplets (“droplets”, *right inlet*) being merged with PCR reagents (*left inlet*) and barcoding beads (*center inlet*) using the Mission Bio Tapestri cartridge. Scale bar = 200 µm. (**C**) Custom microfluidic device design for making tagmenation droplets. Scale bar = 1,000 µm. (**D**) Custom microfluidic device design for merging the tagmentation droplets with PCR reagent and barcoding beads. Scale bar = 1000 µm. Orange reticle indicates region expanded within the right orange frame. Scale bar = 500 µm.

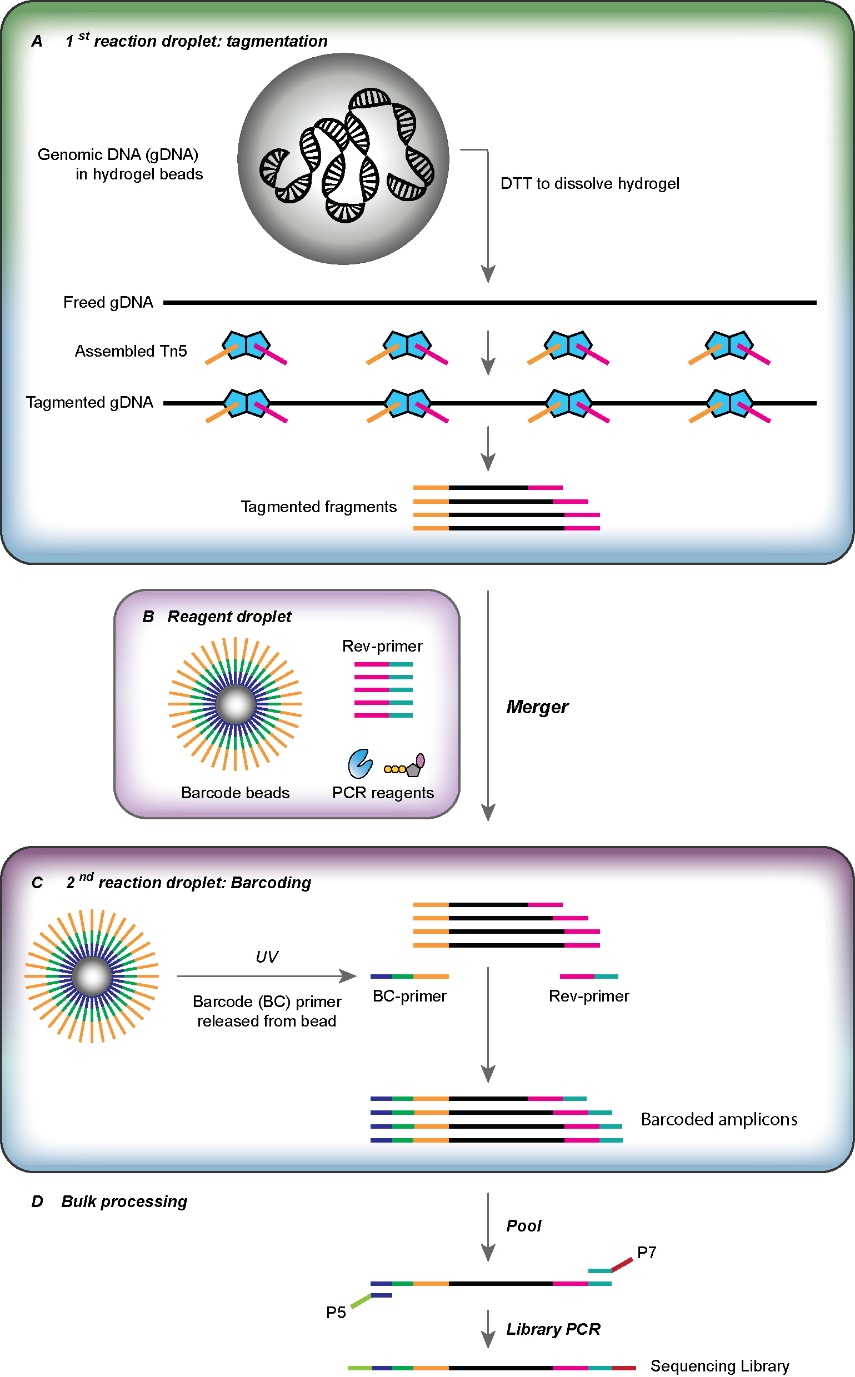

**Fig. S3. EASi-seq molecular biology**.

(**A**) In the 1^st^ reaction droplet, the hydrogel beads are dissolved by dithiothreitol (DTT) to release the genomic DNA (gDNA) and allow for full tagmention by Tn5 transposase. (**B**) The barcode bead carrying barcoding prmer (BC-primer), reverse primer (Rev-primer), and PCR reagents are introduced as an on-demand generated droplet. This reagent droplet is merged with the first reaction droplets. (**C**) Within the formed 2^nd^ reaction droplet, the BC-primers are released from the barcode beads using an ultraviolet (UV) light treatment to break a UV-cleavable linker (IDT code: /iSpPC/). The tagmented DNA fragments are then barcoded and amplified using the BC-primer and Rev-primer in the subsequent droplet digital PCR. (**D**) The droplets, now all containing barcoded amplicons, are broken to allow for pooling of the DNA. A final bulk PCR is then used to introduce the Illumina sequencing adaptors and prepare the sequencing library.

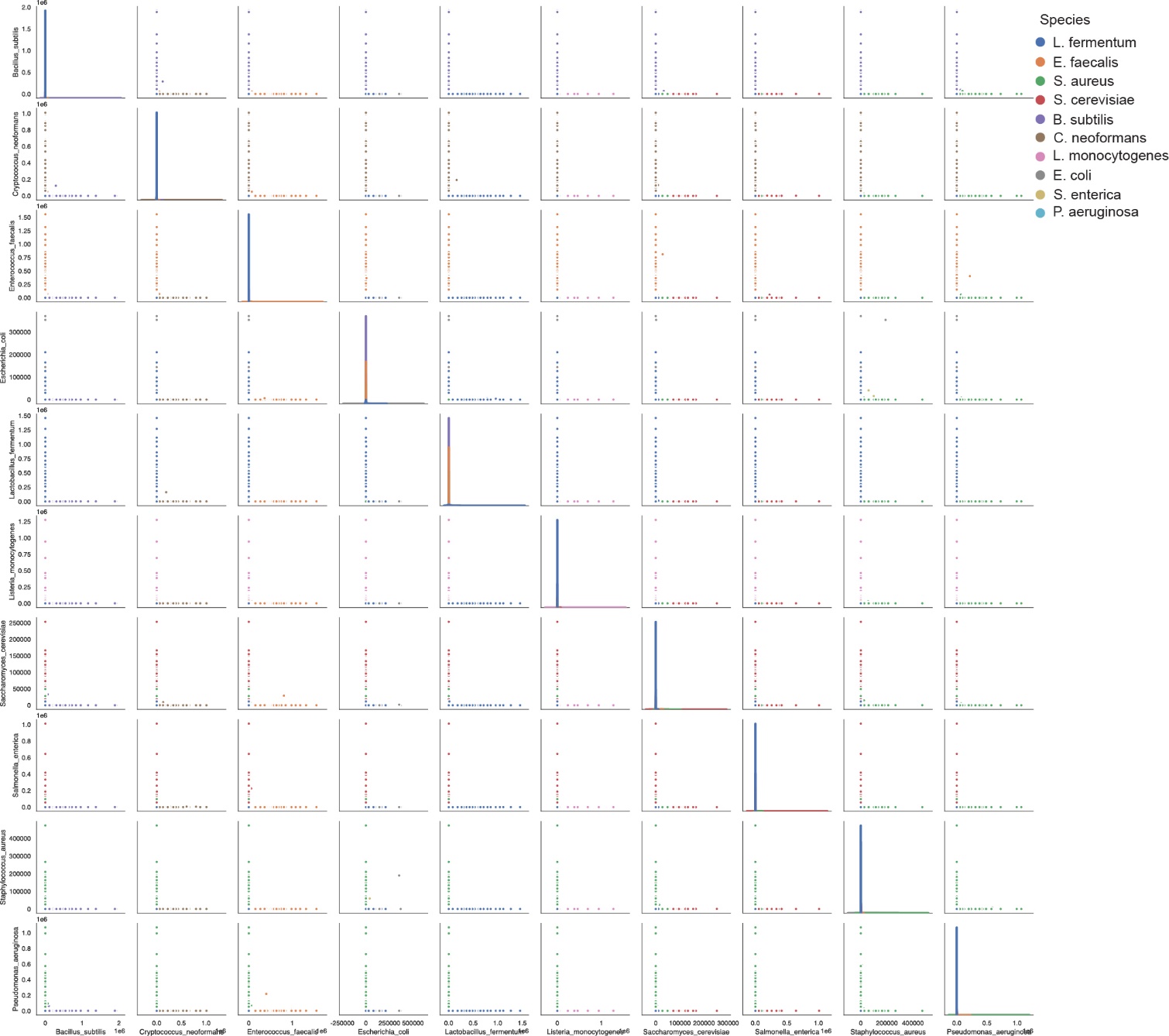

**Fig. S4. Single cell precision achieved across ZymoBIOMICS Microbial Community Standard.** Single cell sequencing barnyard plots, where each dot represents one barcode group and is color-coded by species (mixtures indicated by grey).

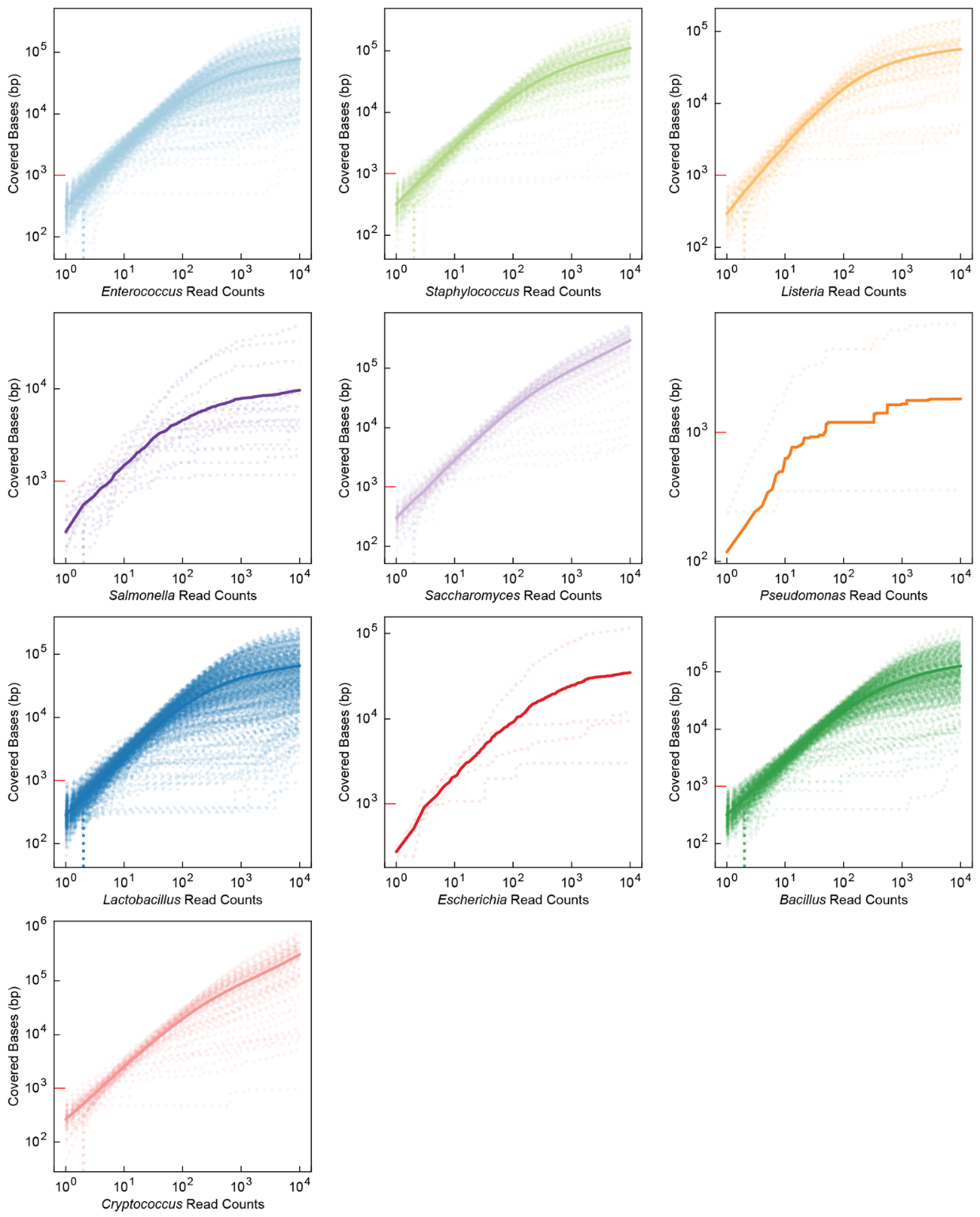

**Fig. S5. Rarefaction analysis of ZymoBIOMICS barcode groups by species.**

For each barcode group of a given species in the ZymoBIOMICS sample, 10,000 reads were randomly sampled from the alignment file (SAM file) of each barcode group and the total coverage was tabulated after each sampling. Dotted lines represent the number of covered bases as a function of sampled read counts. Solid lines represent the averaged coverage across all barcode group of the corresponding species. As a scale reference, the 10^3^ bp marker is colored red in all plots.

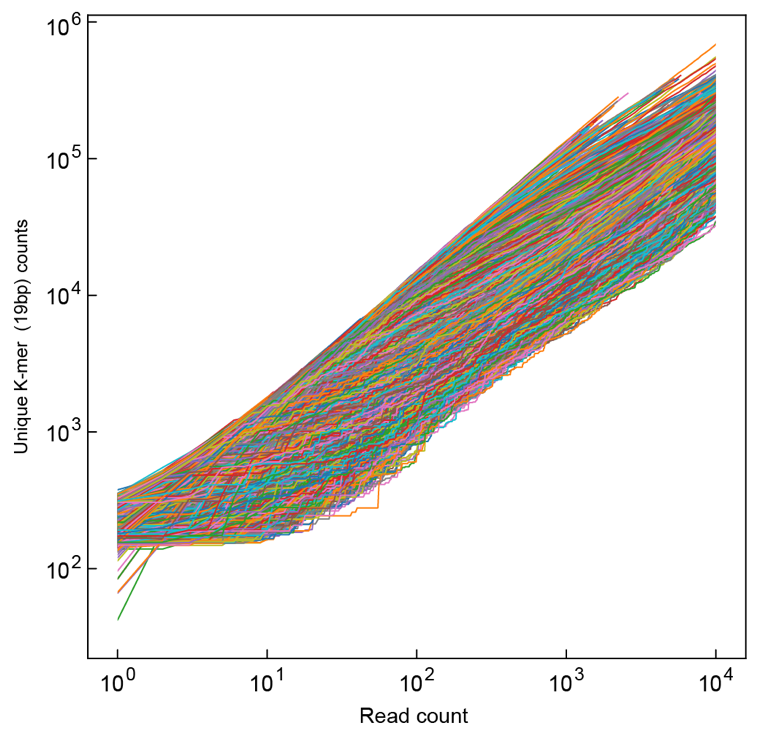

**Fig. S6. Rarefaction analysis of ZymoBIOMICS barcode groups by unique K-mer counting.**

For all barcode groups in the ZymoBIOMICS sample, 10,000 reads were randomly sampled from the raw read file (FASTQ file) and the total number of unique K-mers (19bp) were counted after each sampling. Each line represents one barcode group.

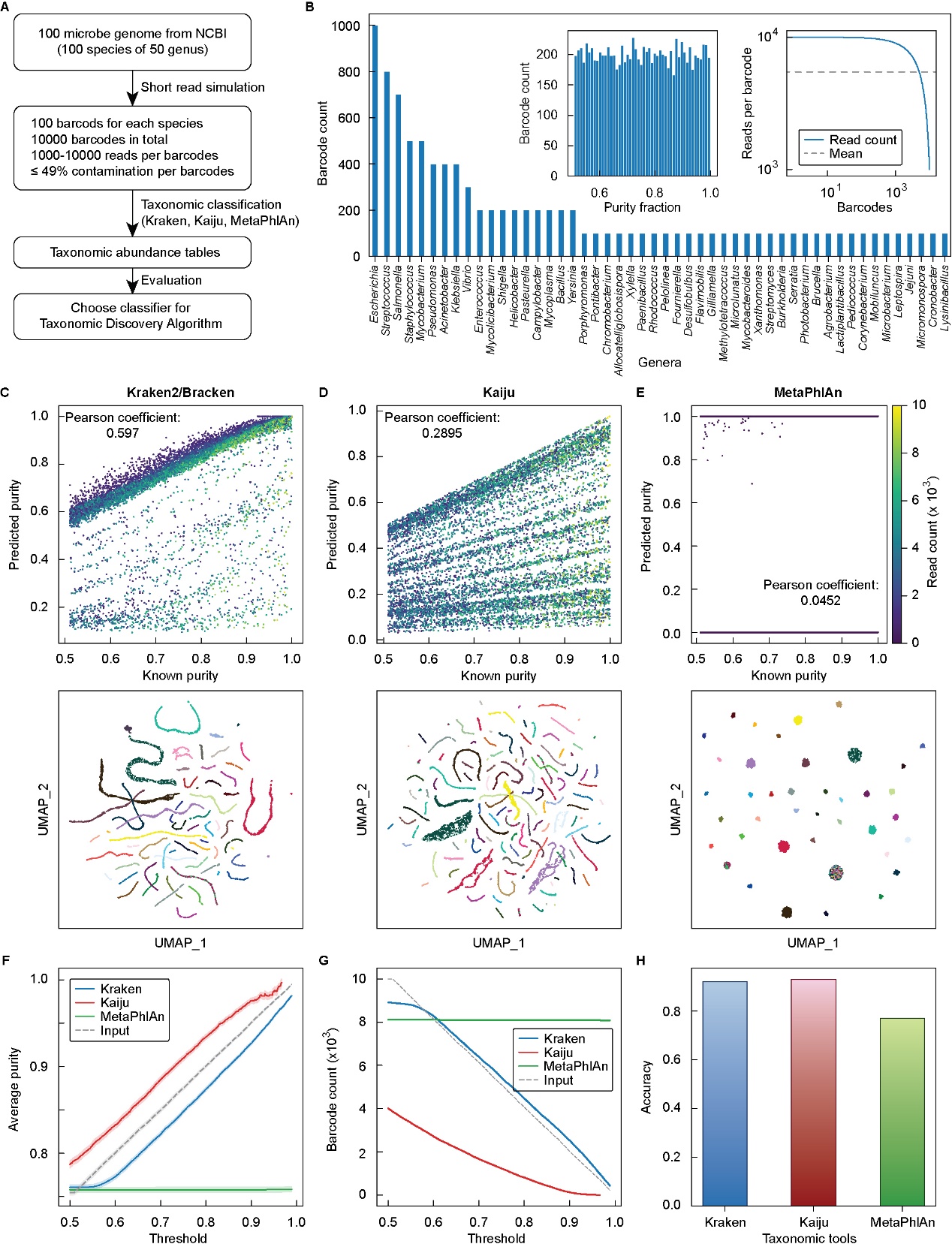

**Fig. S7. Taxonomic classifier selection for taxonomic discovery algorithm (TDA) using simulated single cell sequencing data.**

(**A**) Experimental design of simulation. Genome sequences (FASTA format) of one hundred random chosen species, spanning fifty genera, were downloaded from NCBI. Next, one hundred simulated short read files were generated for each genome. Each file contains up to 49% contaminating reads randomly generated from the other ninety-nine genomes. The read count of each short read file ranged between 1,000 to 10,000. Each short read file (or barcode group) was analyzed using three different taxonomic classifier tools (Kraken2/Bracken, Kaiju, MetaPhlAn). The results were evaluated to find the best taxonomic classifier for barcode group analysis. (**B**) genus distribution of the randomly downloaded genomes. *Left inset*: purity distribution of the 10,000 short read files. *Right inset*: Read counts of the short read files. (**C-E**) Taxonomic classifier analysis for the three tools. *Upper plots*: purity correlation plot (predicted vs known). *Lower plots*: UMAP clustering based on generated taxonomic abundance tables for simulated barcode groups, with color coding by genus. (**F**) Average purity calculated after data filtering with thresholds ranging from 0.5 to 0.99. Shaded areas indicate the 95% confidence intervals. (**G**) Number of barcodes passing filter when using thresholds ranging from 0.5 to 0.99. (**H**) Accuracy of the UMAP clustering based on taxonomical assignment.

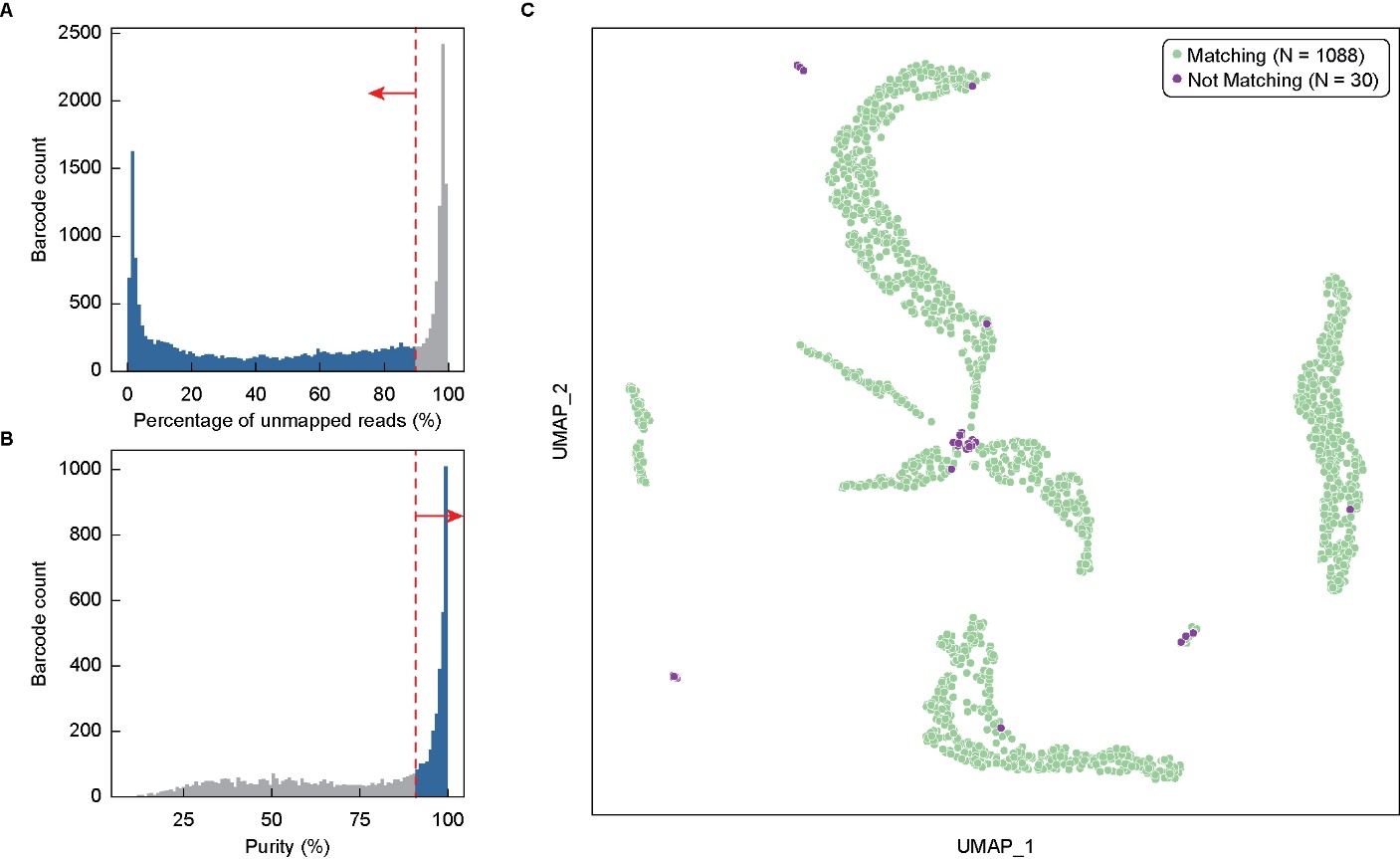

**Fig. S8. ZymoBIOMICS single cell sequencing data filtering with TDA and UMAP evaluation**.

(**A**,**B**) Barcode group filtering with percentage of unmapped reads (**A**) and genus level purity (**B**). (**C**) UMAP clustering by the taxonomic discovery algorithm (TDA) using the estimated genus abundance of barcode groups that passed filtering. The UMAP plot highlights the difference between reference genome-based identification and Kraken2-based identification (as reported in **Fig. 2G**).

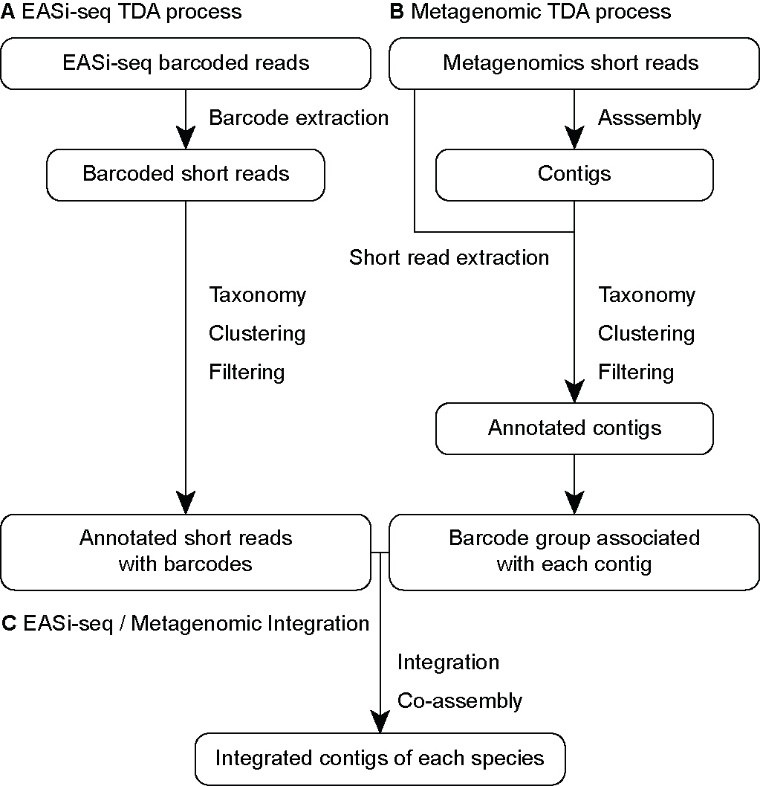

**Fig. S9. Flowchart of EASi-seq and metagenomic data integration process.** (**A**) The barcode sequences are extracted from EASi-seq raw reads and used to assign reads to barcode groups. The taxonomy abundance of each barcode group is estimated before clustering by the taxonomy discovery algorithm (TDA). (**B**) Concurrently, the metagenomic reads are assembled into contigs, and the raw reads are mapped back to the contigs to identify the raw short reads associated with each contig. The reads associated with a given contig are treated as a barcode group and processed with TDA to generate annotated contigs. (**C**)The EASi-seq barcode groups and contig barcode groups are integrated and clustered. All the reads from the barcodes in the same cluster are pooled and co-assembled.

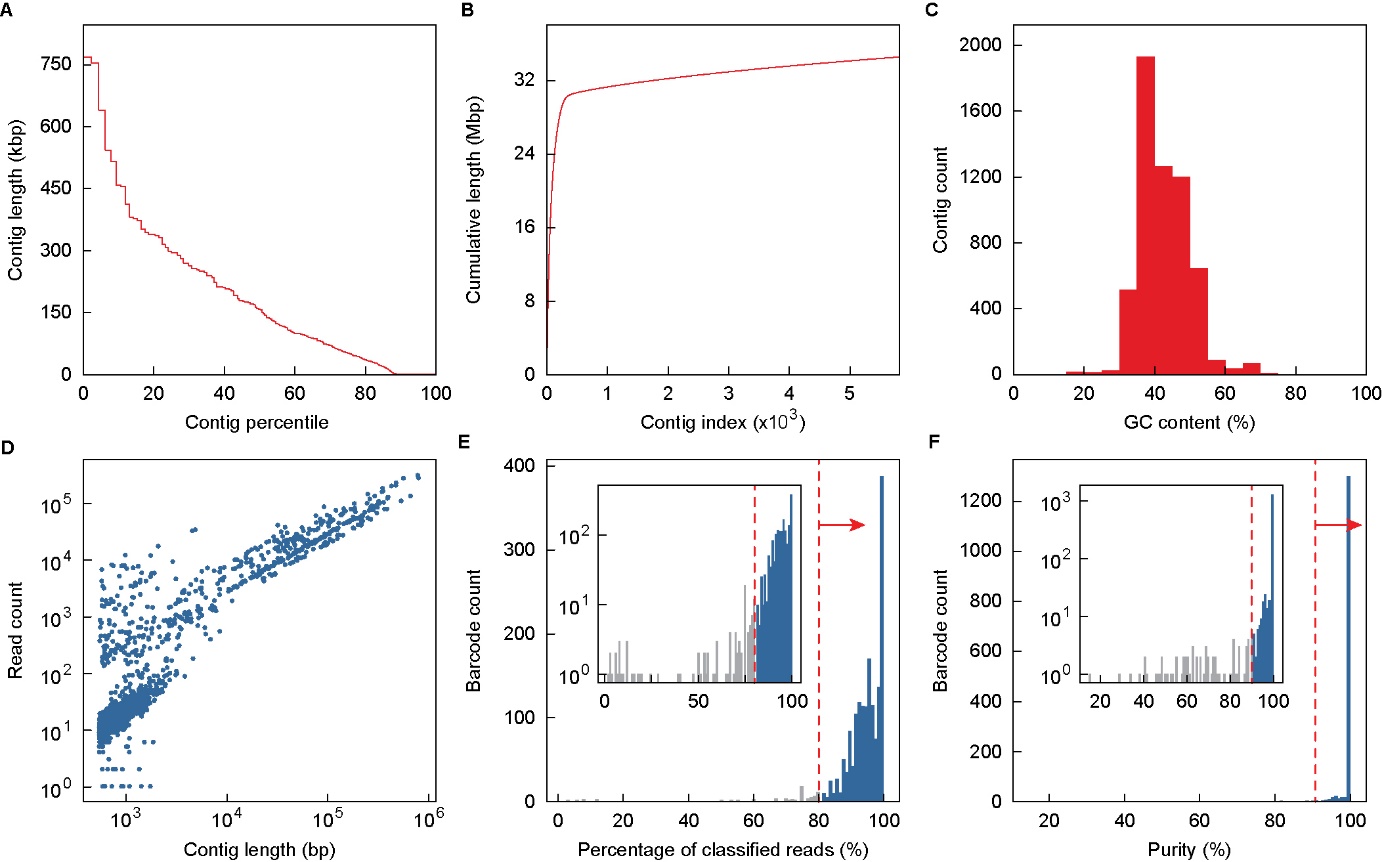

**Fig. S10. Quast evaluation of the ZymoBIOMICS metagenomic assembly and contig filtering.**

(**A-C**) The contig length distribution (**A**), accumulative contig length (**B**), and GC content (**C**) of the ZymoBIOMICS metagenomic assembly. The metagenomic shotgun reads of the ZymoBIOMICS community were assembled into contigs before the quality was evaluated by Quast. Each contig was treated as a barcode group and pair end reads associated to each contig were extracted. (**D**) Correlation analysis between a contig’s length and the number of associated reads found after alignment of the raw read data. Associated reads for each contig are treated as a barcode group in subsequent analysis. (**E**-**F**) Filtering of contig barcode groups by percentage of classified reads (**E**) and then predicted genus level purity (**F**). Contig barcode groups are processed by the taxonomic discovery algorithm (TDA). Contig barcode groups with less than 80% reads that are classified by Kraken2 were removed (**E**). Purity was calculated as percentage of reads with a TDA classification same as the contig barcode group’s dominate taxa. Contig barcode groups with less than 90% purity were removed (**F**). Remaining barcode groups were integrated with the single cell barcodes (as shown in **Fig. 2H**).

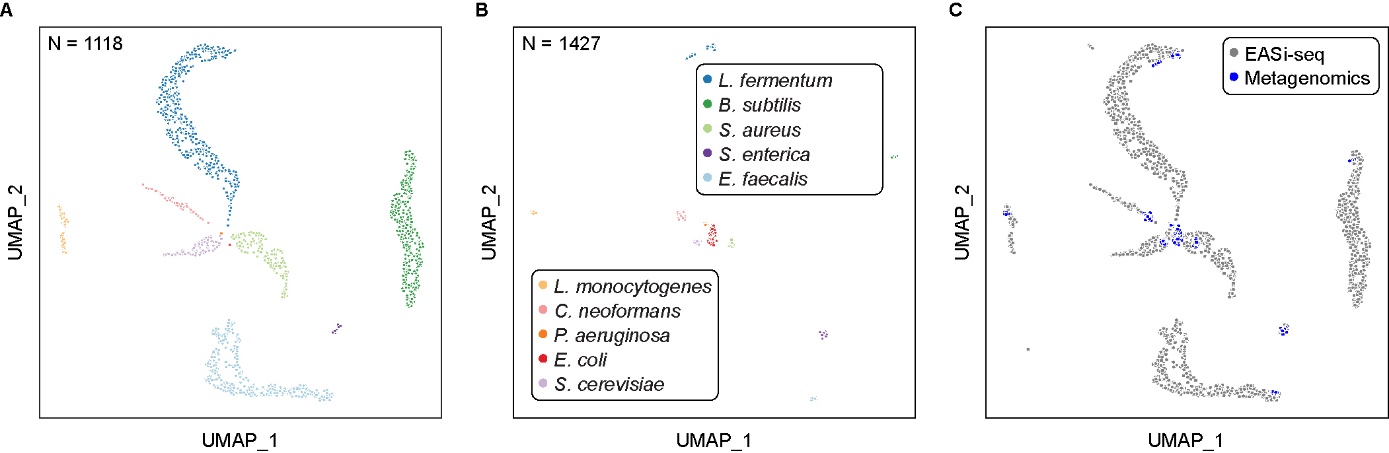

**Fig. S11. EASi-seq and metagenomic data integration for ZymoBIOMICS sample.**

(**A**) EASi-seq UMAP clustering by TDA with colors indicating genus annotation, reproduced from main text (**Fig. 2G**). (**B**) Metagenomic contigs UMAP clustering by TDA with colors indicating genus annotation. (**C**) Integrated UMAP of both EASi-seq barcodes and Metagenomic contigs showing overlap and augmentation on top of (**A**), reproduced from main text (**Fig. 2H**).

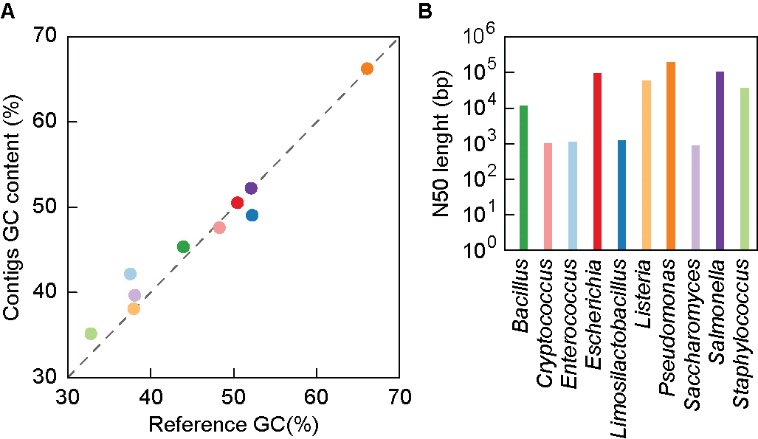

**Fig. S12. GC content and N50 analysis of assembled contigs from ZymoBIOMICS TDA clusters.**

(**A**) GC content comparison between reference genome and assembled contigs. (**B**) N50 of assembled contigs grouped together based on TDA.

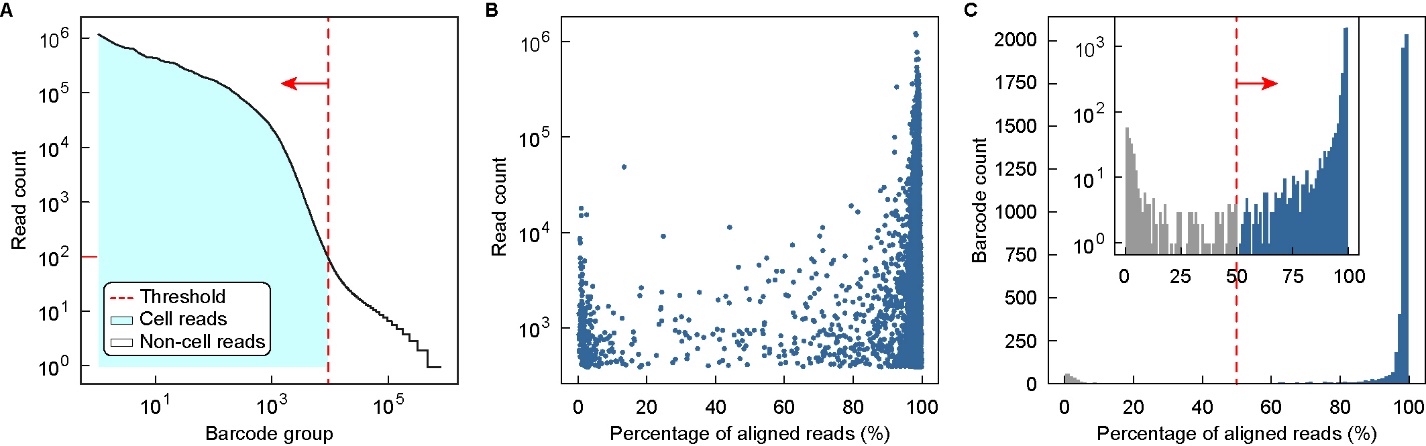

**Fig. S13. EASi-seq data filtering for synthetic community of twenty-two *Eggerthella lenta* strains**. (**A**) Barcode groups were first filtered by read count, using 100 reads as the cutoff for retention. (**B**) Correlation analysis between barcode groups’ read count and alignment percentage after mapping to the reference genome. (**C**) Further filtering removed barcode groups with less than 50% alignment rates.

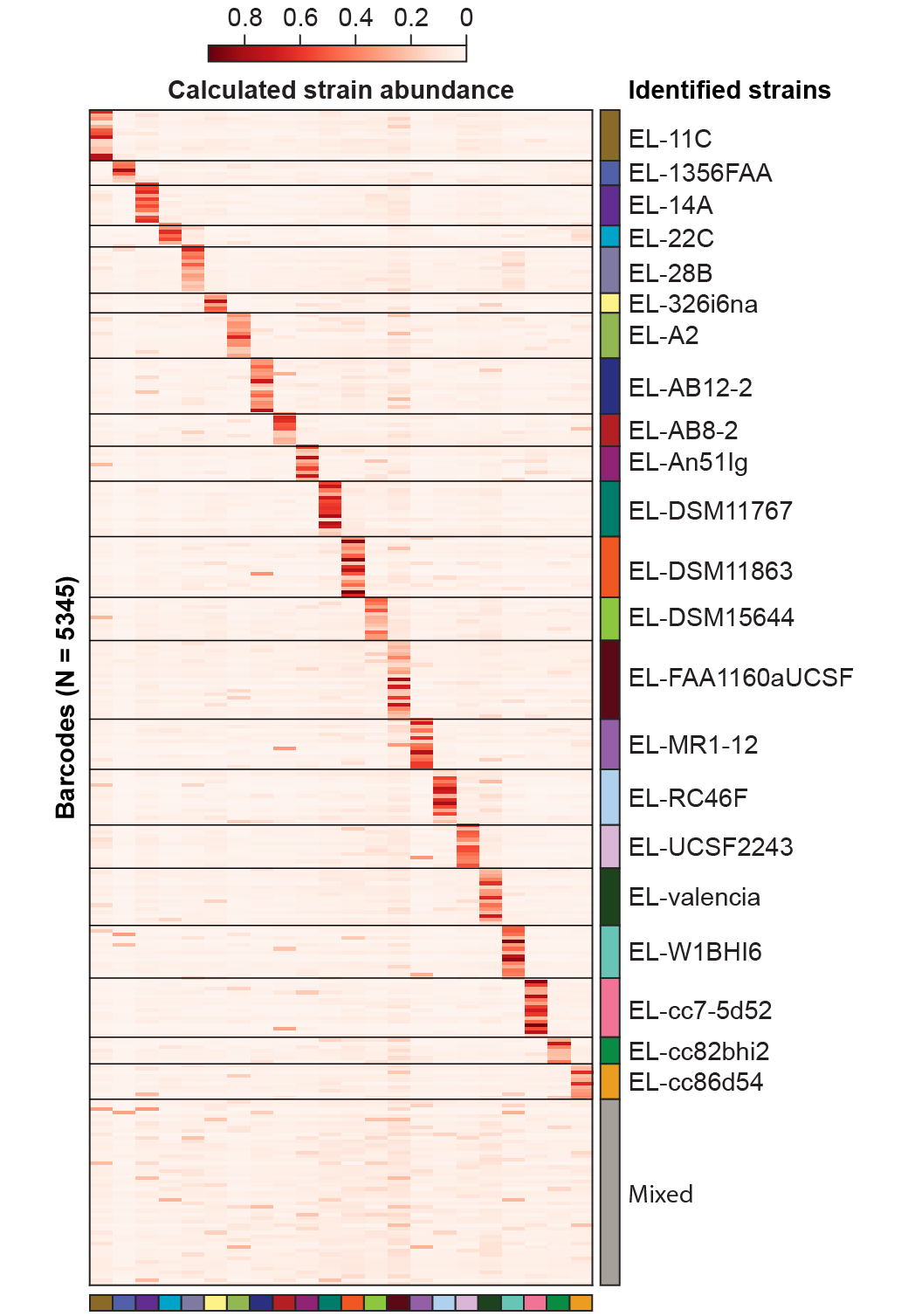

**Fig. S14. EASi-seq barcode group annotation for *E. lenta* synthetic community**.

Heatmap of strain abundance in each barcode group, grouped by identified strain. Reads in each barcode group (*rows*) are mapped to the twenty-two reference genomes (*columns*), with the abundances of each strain in a given barcode calculated by Bayesian estimation. Strain coding corresponds to colors used in main text (**Fig. 3**). Gray indicates mixed/unresolved barcodes.

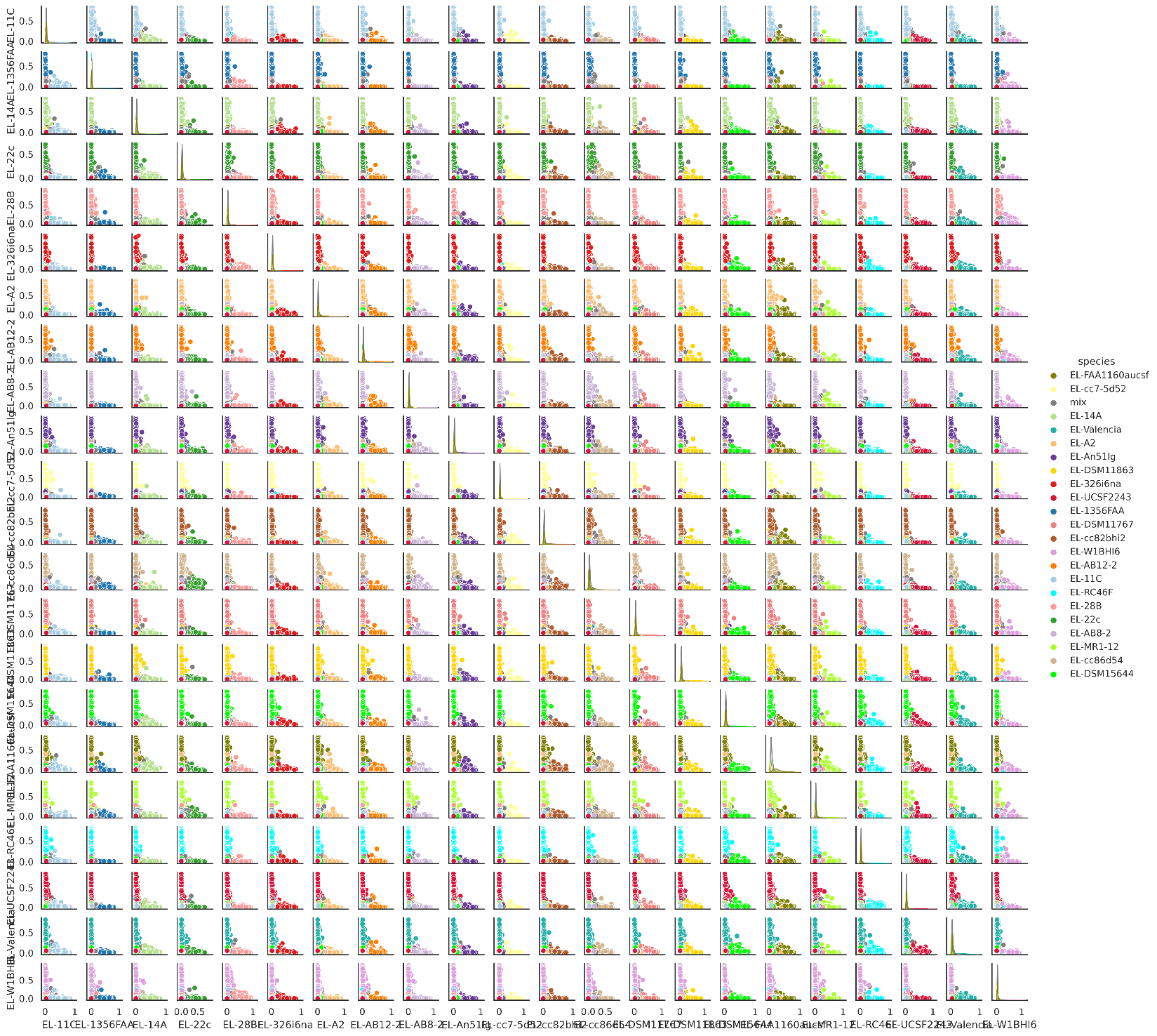

**Fig. S15. Mixing analysis based on estimated abundance of E. lenta strains in each barcode**.

Barnyard plots show the estimated pair-wise abundance of corresponding *E. lenta* strains. Each dot represents one barcode group and is color coded by identified strain.

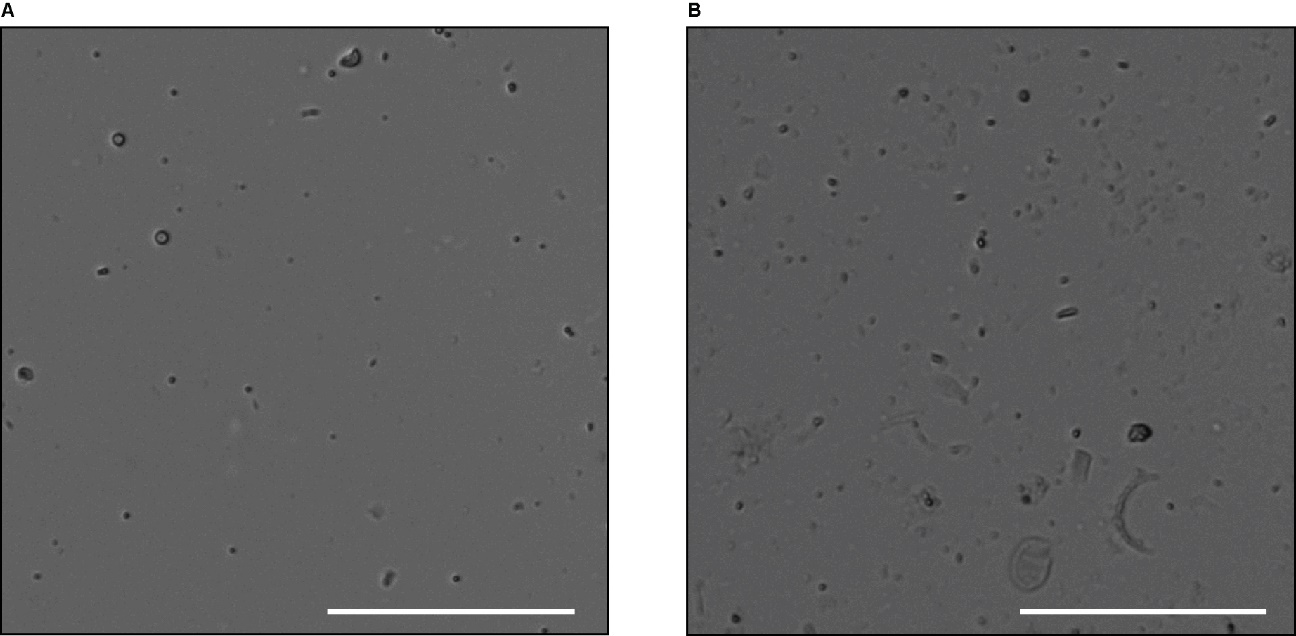

**Fig. S16. Isolated microbe suspensions used for EASi-seq.** (**A**) Cells isolated from human fecal sample by density centrifugation. (**B**) Cells isolated from Ocean Beach sea water by filtration. Scale bar = 100 µm.

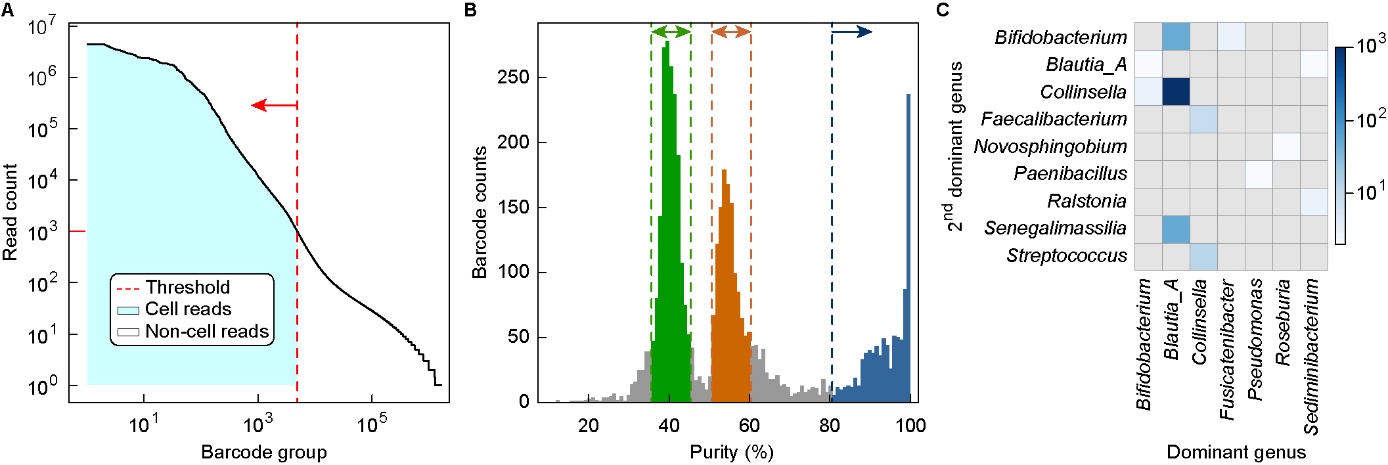

**Fig. S17. Cell aggregation interactions in frozen human fecal sample.**

(**A**) Barcode groups were first filtered by read count, using 1000 reads as the cutoff for retention. (**B**) The taxonomical purity distribution showed two peaks centered at 40% (*green*) and 55% (*orange*), repectively indicating barcode groups with three or two cells. Purities over 80% (*blue*) are for barcode groups containing only one cell. (**C**) Cell-cell interaction count for barcode groups within the 50%-60% purity interval.

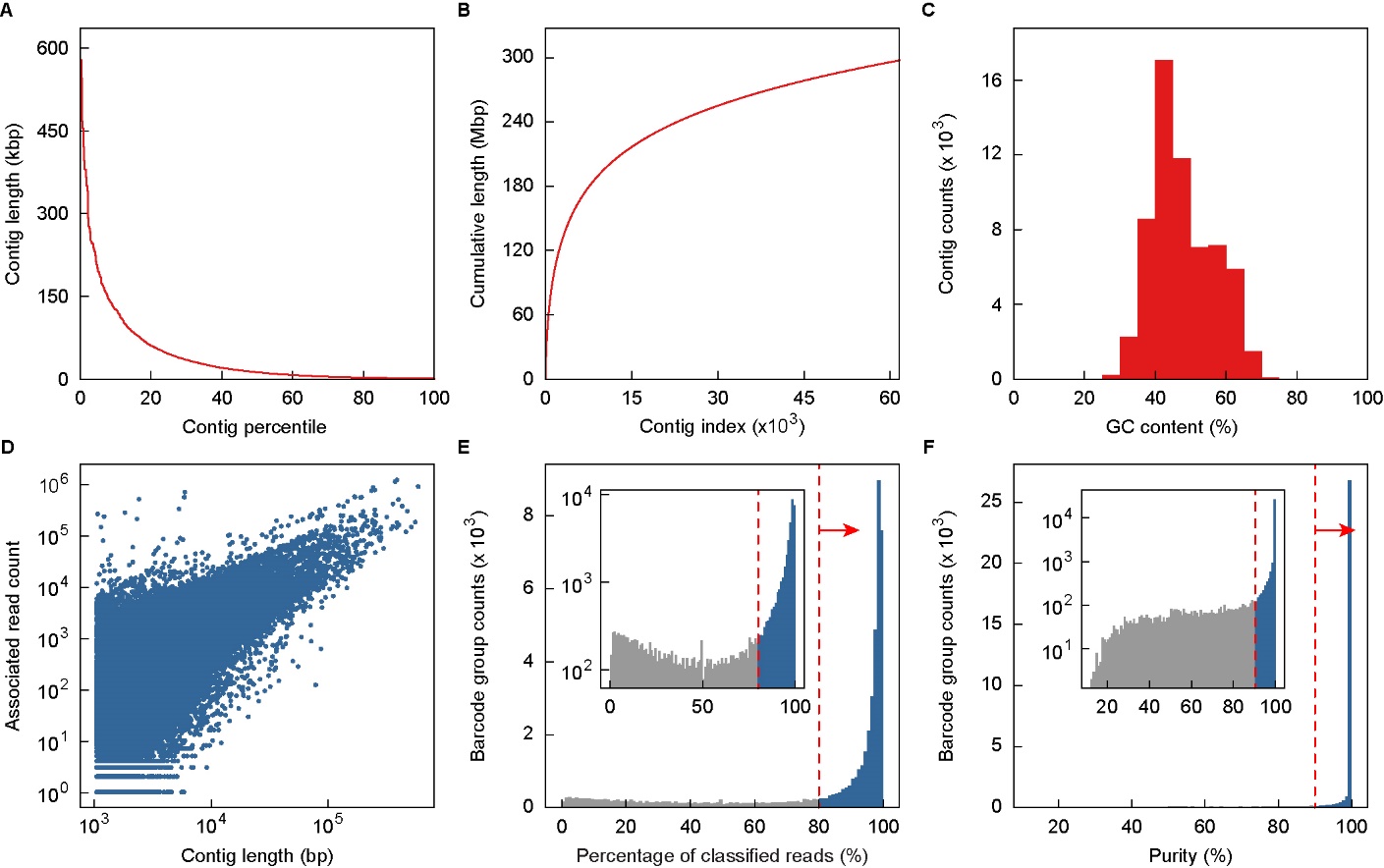

**Fig. S18. Quast evaluation of human microbiome metagenomic assembly and contig filtering.** (**A-C**) Human human microbiome metagenomic assembly statistics of contig length distribution (**A**), accumulative contig length (**B**), and GC content (**C**). (**D**) Correlation analysis between a contig’s length and the number of associated reads found after alignment of the raw read data. Associated reads for each contig are treated as a barcode group in subsequent analysis. (**E**) Contig barcode group are first filtered by percentage of reads classified by TDA. (**F**) Contig barcode groups are then filtered by purity, calculated as percentage of reads with a TDA classification same as the contig barcode group’s dominate taxa. The remaining contig read groups were integrated with the single cell barcodes used in for downstream analysis.

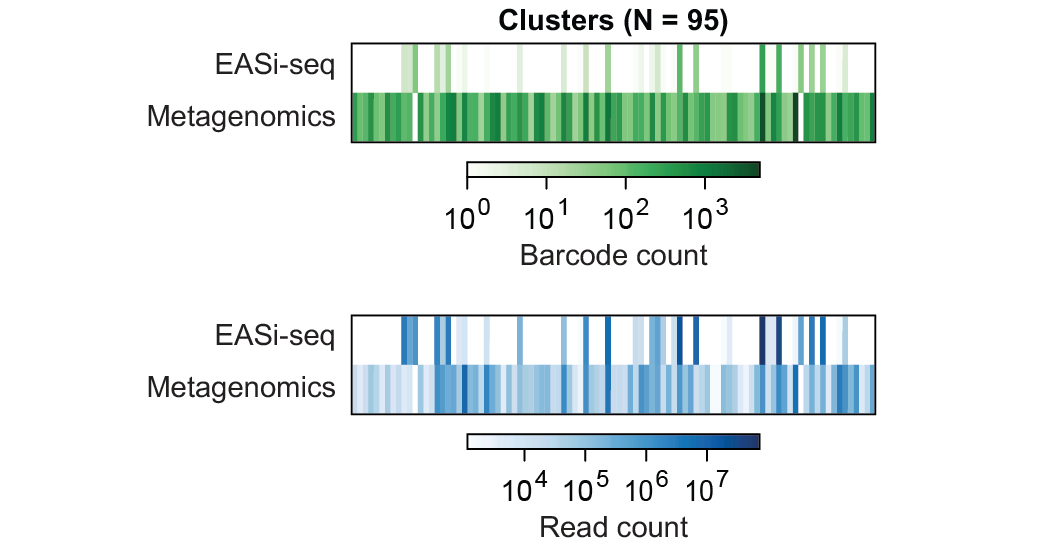

**Fig. S19. Human microbiome barcode and read count analysis per TDA cluster.**

Barcode counts (*green*) and associated read counts (*blue*) from filtered EASi-Seq and metagenomic data based on TDA clusters.

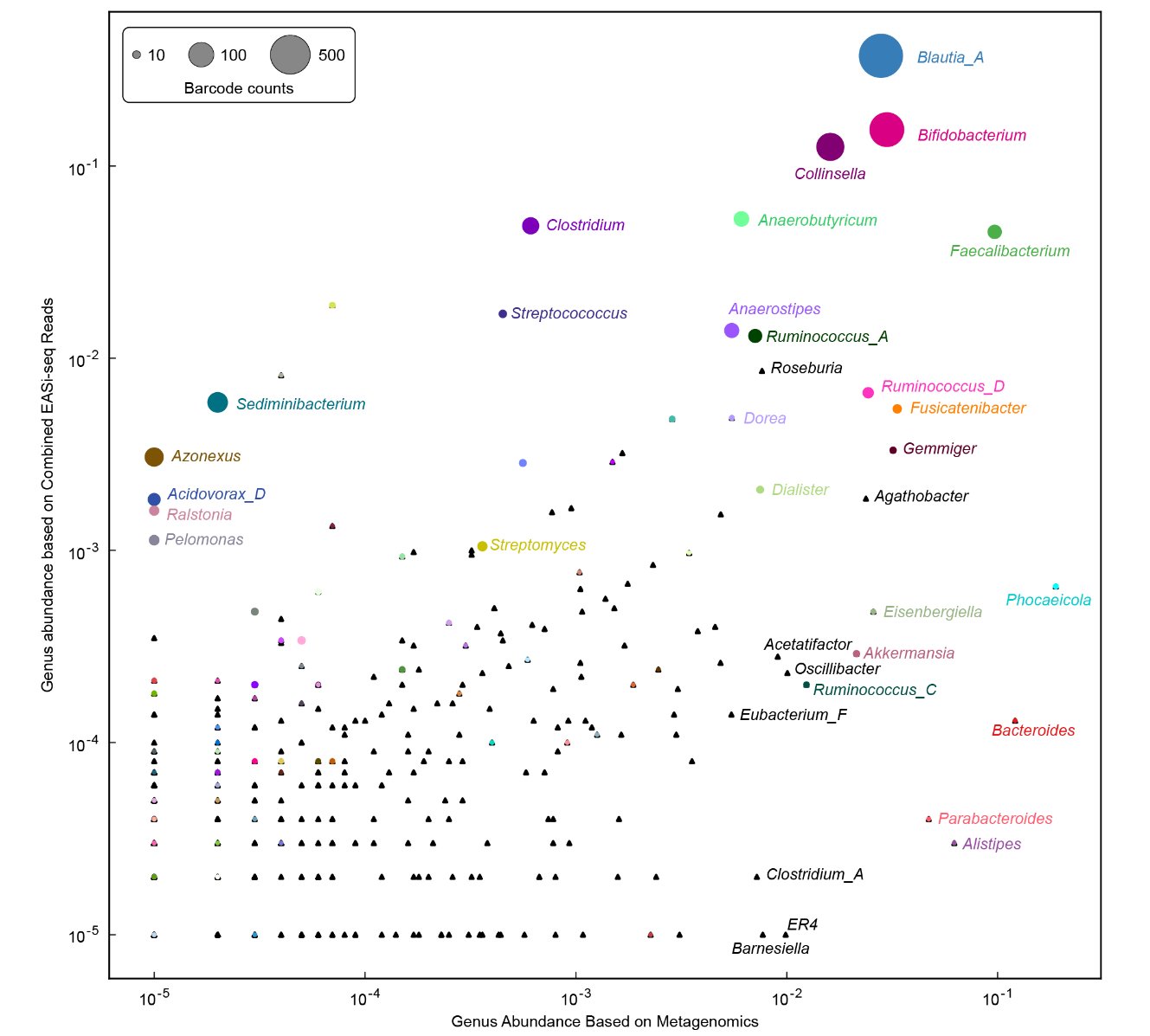

**Fig. S20. Comparison between metagenomic and EASi-seq outputs for human microbiome sample.**

Scatter plot shows the relative abundance of genus found by Kraken2 in the metagenomics sequencing data and the combined EASi-seq barcodes groups (includes all barcodes before filtering). All genera discovered by TDA are shown in circles and scaled by barcode counts, with the rest genera without any associated barcode shown in triangles. Labeled genera exist at either >0.5% abundance in the metagenomics data or with >5 barcode groups in the EASi-seq data.

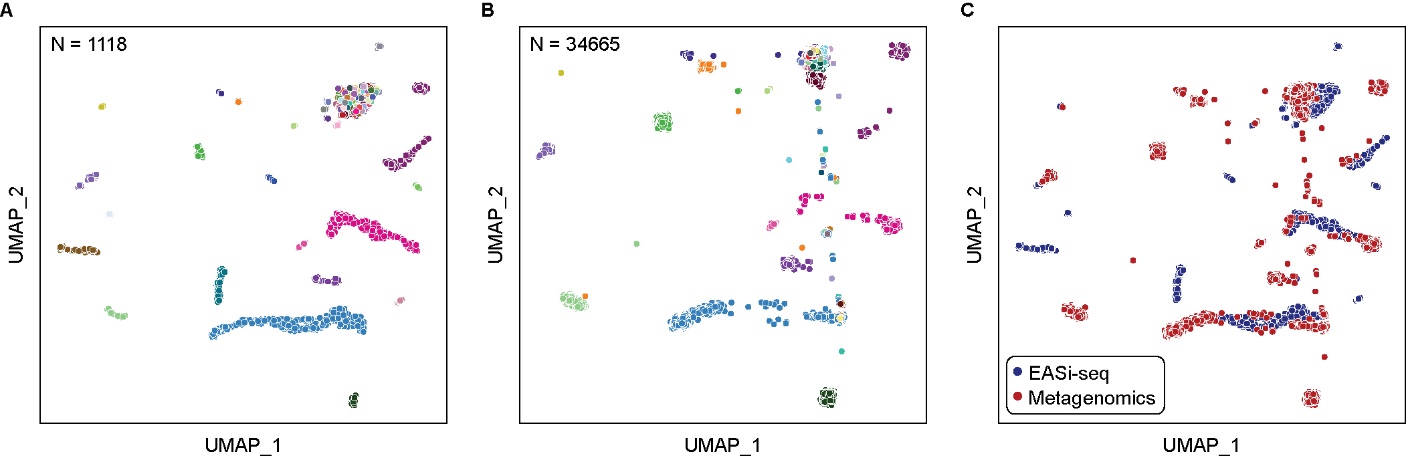

**Fig. S21. EASi-seq and metagenomic data integration for human microbiome sample.** (**A**) UMAP clustering of only EASi-seq data based on TDA, color coded by genus annotation. (**B**) UMAP of clustering metagenomic contig data by TDA, color coded by genus annotation. (**C**) Integrated UMAP using both EASi-seq barcodes and Metagenomic contigs, color coded by data point source.

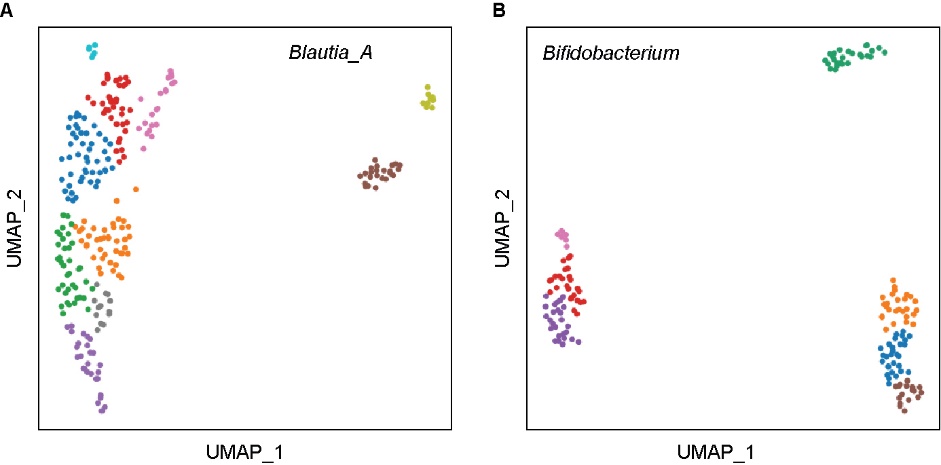

**Fig. S22. Subclustering of EASI-seq barcodes in the top two TDA clusters.**

(**A**) UMAP corresponding to EASi-seq barcode groups in TDA cluster for *Blautia_A*. (**B**) UMAP corresponding to EASi-seq barcode groups in TDA cluster for *Bifidobacterium*. All TDA clusters based on Kraken2 species level estimates.

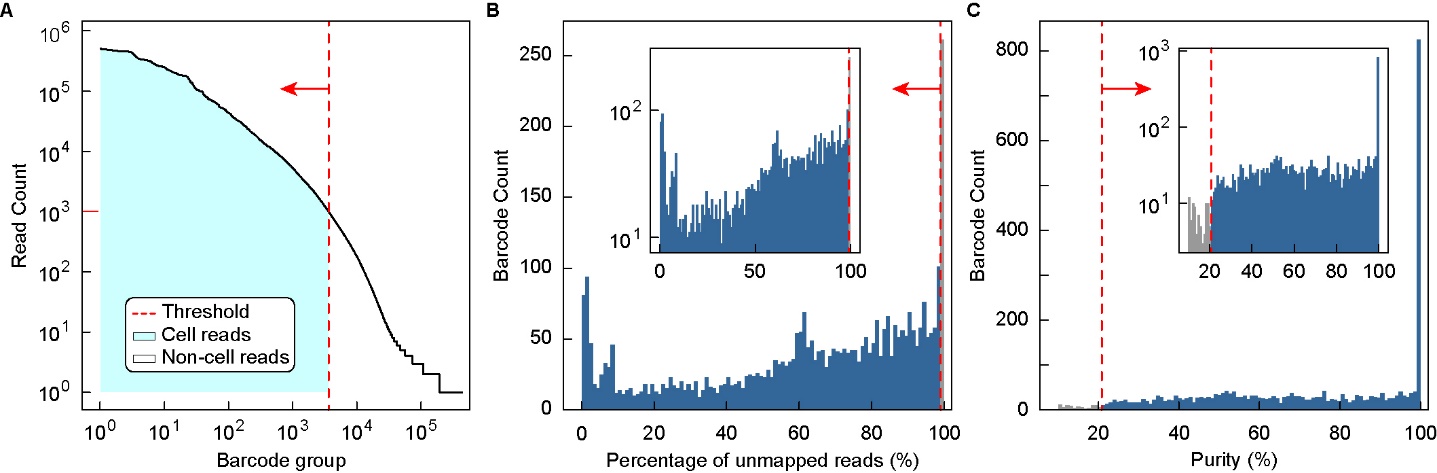

**Fig. S23. Ocean Beach sample data filtering.**

(**A**) Barcode groups were filtered by read count, using 1000 reads as the cutoff for retention. (**B**) Barcode group are filtered by percentage of reads that are not classified by TDA (barcode groups with more than 99% reads that are not mapped to the Kmer database were removed). (**C**) Barcode groups are then filtered by purity, calculated as percentage of reads with a TDA classification same as the barcode group’s dominate taxa.

**Table S1:** Evaluated commercial platforms to develop single microbe sequencing.

| **Company** | **Platform** | **Single cell operation mechanism** | **Website** |
| --- | --- | --- | --- |
| 10X | Chromium | Droplet microfluidics | <https://www.10xgenomics.com/instruments/chromium-controller> |
| Bio-Rad | ddSEQ | Droplet microfluidics | <https://www.bio-rad.com/en-us/life-science/digital-pcr/single-cell-sample-preparation-for-ngs/ddseq-single-cell-isolator> |
| BD | Rhapsody | Microwell array | <https://go.bd.com/bd-rhapsody.html> |
| Mission Bio | Tapestri | 2 step droplet microfluidics | <https://missionbio.com/products/platform/> |
| Dolomite Bio | Nadia | Droplet microfluidics | <https://www.dolomite-bio.com/product/nadia-instrument/> |
| Celsee | Genesis | microwell | <https://www.bio-rad.com/en-us/category/circulating-tumor-cell-ctc-enrichment-enumeration?ID=3b7d67be-8213-ed2f-0b0a-1ee1f456f801> |
| Fluent Bio | PIPseq | Pre-templated Instant Partitions | <https://www.fluentbio.com/products/pipseq-3-single-cell-rna-kit/> |

**Table S2-S21 description**

**Table S2**: Primer sequences used in this work.

**Table S3**: ZymoBIOMICS Microbial Community Standards species:

**Table S4**: ZymoBIOMICS Microbial Community Standards EASi-seq barcode reference genome based analysis: The rows are barcode groups. The first 4 columns are the summary of reference alignment, including totally mapped read count, read count that mapped to the dominant species, purity, and dominant species. The other columns are divided into 4 groups (10 columns each), which represent the mapped read counts to 10 individual species(readCount@species name), covered bases in 10 individual species (covBases@species name), percentage coverages of 10 individual species (CovBases@species name), and mean depth of 10 individual species (meanDepth@specie name).

**Table S5**: ZymoBIOMICS Microbial Community Standards EASi-seq barcode Taxonomic Discovery Algorithm (TDA) analysis.

**Table S6**: ZymoBIOMICS Microbial Community Standards metagenomic generated contigs Taxonomic Discovery Algorithm (TDA) analysis.

**Table S7**: List of the twenty-two strains of E. lenta included in the synthetic community.

**Table S8**: E. lenta EASi-seq barcode strain identification. The rows are barcode groups. The first two columns (Read_Cnt and mapping_rate) are the total read count and overall mapping percentage of the reads to the reference genome. The last column is the strain annotation of the corresponding barcode group. The other 22 columns are the estimated abundance of each strain.

**Table S9**: Human microbiome EASi-seq barcode Taxonomic Discovery Algorithm (TDA) analysis.

**Table S10**: Human microbiome metagenomic generated contigs Taxonomic Discovery Algorithm (TDA) analysis.

**Table S11**: Human microbiome cluster statistics. The barcode group count, metagenomic contig count, total read count from single cell and total read from metagenomic contigs of each genus clusters.

**Table S12**: Human microbiome EASi-seq barcode antibiotic resistance gene discovery. The rows are genera clusters and the columns are antibiotic resistant genes. Each value in the table represents the relative abundance of antibiotic resistant gene (antibiotic resistance gene read count per Million reads).

**Table S13**: Human microbiome EASi-seq barcode metabolism pathway analysis. The columns are genus clusters. The first 4 columns (pathway_name, function, class, super_class) are the name, function, class and super class of each pathway. The values in the table represent the pathway abundance in the corresponding genus clusters.

**Table S14**: Costal sea water microbiome EASi-seq barcode Taxonomic Discovery Algorithm (TDA) analysis. The columns are barcode groups. The first 4 columns (percent_covered, read_count_covered, read_assigned, total_reads) represent the percentage of reads that are mapped to the k-mer database, count of reads that mapped to the k-mer database, and the total read count in the barcode group. The barcode groups are filtered (percent_covered > 1%). The next columns with genus names starting with ‘g__’ are Kraken/Bracken estimated genus abundance of the corresponding barcode group. The barcode group were further filtered by the abundance of the dominant genus (> 80%). The last few columns are the UMAP clustering and the taxa annotation. (missing value indicate the barcode group did not pass the filter).

**Table S15**: Coastal sea water microbiome EASi-seq barcode clustering statistics. The UMAP coordinate, read counts, barcode count, NCBI_id, and taxa annotation at various level (Domain, Phylum, Class, Order, Family, Genus) of each cluster identified in the EASi-seq barcodes.

**Table S16**: Halioglobus single cell gene analysis. The columns are Halioglobus barcodes, and the rows are genes identified by HUMANN. Each value in the table represents abundance (RPKs, read count per thousand reads) of the gene in the corresponding barcode group.

**Table S17**: 100 strains for TDA testing (species, NCBI_ID).

**Table S18**: 10000 simulated barcode group from the 100 strains (barcode name, purity, total reads).

**Table S19**: Simulation data analysis with Kraken2/Bracken. The rows are barcode groups. The first two columns (Input purity and predicted purity) are the purity of reads in barcode group by design and by Kraken2/Bracken prediction. The third to fifth columns (leiden, celltype, genera_input) are the result of UMAP clustering. Leiden represent the clusters identified by UMAP. Celltype represents the predicted genus and the genera_input are the genus used to generate the simulation barcode group. The rest of the columns are the Kraken2/Bracken generated abundance of corresponding genus listed as the column names.

**Table S20**: Simulation data analysis with MetaPhlAn. The rows are barcode groups. The first two columns (Input purity and predicted purity) are the purity of reads in barcode group by design and by MetaPhlAn prediction. The third to fifth columns (leiden, celltype, genera_input) are the result of UMAP clustering. Leiden represent the clusters identified by UMAP. Celltype represents the predicted genus and the genera_input is the genus used to generate the simulation barcode group. The rest of the columns are the MetaPhlAn generated relative abundance (percentage) of corresponding genus listed as the column names.

**Table S21**: Simulation data analysis with Kaiju. The rows are barcode groups. The first two columns (Input purity and predicted purity) are the purity of reads in barcode group by design and by Kaiju prediction. The third to fifth columns (leiden, celltype, genera_input) are the result of UMAP clustering. Leiden represent the clusters identified by UMAP. Celltype represents the predicted genus and the genera_input is the genus used to generate the simulation barcode group. The rest of the columns are the Kaiju generated relative abundance (percentage) of corresponding genera listed as the column names.
